# Molecular and spatial profiling of the paraventricular nucleus of the thalamus

**DOI:** 10.1101/2022.07.19.500649

**Authors:** Claire Gao, Chiraag A. Gohel, Yan Leng, David Goldman, Ariel J. Levine, Mario A. Penzo

**Author notes:** Correspondence Claire Gao, B.A.;, Mario A. Penzo, Ph.D. These authors contributed equally (Chiraag A. Gohel, Yan Leng).

## Abstract

The paraventricular nucleus of the thalamus (PVT) is known to regulate various cognitive and behavioral processes. However, while functional diversity among PVT circuits has often been linked to cellular differences, the molecular identity and spatial distribution of PVT cell types remains unclear. To address this gap, here we used single nucleus RNA sequencing (snRNA-seq) and identified five molecularly distinct PVT neuronal subtypes. Additionally, multiplex fluorescent *in situ hybridization* of top marker genes revealed that PVT subtypes are organized by a combination of previously unidentified molecular gradients. Lastly, comparing our dataset with a recently published single-cell sequencing atlas of thalamus yielded novel insight into the PVT’s connectivity with cortex, including unexpected innervation of auditory and visual areas. This comparison also revealed that our data contains a largely non-overlapping transcriptomic map of multiple midline thalamic nuclei. Collectively, our findings uncover previously unknown features of the molecular diversity and anatomical organization of the PVT and provide a valuable resource for future investigations.

## INTRODUCTION

Recent models describe the PVT as a midline thalamic structure that integrates cortical, hypothalamic, and brainstem signals to drive adaptive behavioral strategies amid challenging situations [1–5]. Consistent with this model, neuronal activity in the PVT is sensitive to both interoceptive and exteroceptive salient signals including hunger, reward, punishment, fear, and environmental cues that predict either positive and/or negative outcomes. In turn, the PVT plays a critical role in the signaling of emotional and motivational states largely via projections to the cortex, amygdala, and ventral striatum [7–20]. Given the PVT’s involvement in such a diverse array of functions, there is growing interest in determining how this structure is organized into functional subnetworks, particularly since broad manipulations of the PVT have often yielded disparate results [6, 7]. For instance, while recent reports support the existence of a causal relationship between increased neuronal activity in the PVT and wakefulness, activation of a subpopulation of PVT neurons decreases wakefulness and promotes NREM sleep [8–10]. Similar functional heterogeneity has been observed in the PVT’s contributions to appetitive behaviors, with some studies reporting that lesions or pharmacological inactivation of the PVT increase food intake [11, 12], while others show that direct or indirect activation of the PVT increases food-seeking behaviors [13–15].

Evidence suggests that the seemingly opposing roles of the PVT might be attributable to network differences that distribute along the rostro-caudal axis of the PVT [6, 16]. Accordingly, the anterior and posterior subregions of the PVT (aPVT and pPVT, respectively) differentially innervate the amygdala, bed nucleus of the stria terminalis (BNST), hypothalamus, prefrontal cortex (PFC), ventral subiculum, and nucleus accumbens (NAc) and have been tied to different functions [17–20]. Although this broad classification into aPVT and pPVT subregions has aided advances in our understanding of the functional organization of the PVT, systematic dissections of the local organization of PVT subnetworks are currently lacking [6, 7, 16]. Importantly, consistent with the notion that molecular identity can delineate cell types with different anatomical, functional, and electrophysiological properties [21–26], studies have recently identified functional and anatomical differences that tie onto genetically defined subpopulations of PVT neurons [8, 10, 27, 28]. From this perspective, transcriptional profiling of the PVT could lead to valuable insights into the functional heterogeneity of this thalamic structure [7].

In this study, we employ high throughput single-nucleus RNA sequencing and multiplex fluorescent *in situ* hybridization (ISH) labeling to identify five PVT subtypes that segregate differentially across the antero-posterior, medio-lateral, and dorso-ventral axes of the PVT. Our data highlights novel features about the molecular organization of the PVT, thereby offering a glimpse into how genetic diversity may tie into the various functions associated with this region of thalamus. In addition, by performing reciprocal principal component analysis (RPCA)-based integration and comparative analysis with the ThalamoSeq atlas, we find that PVT subtypes differentially innervate cortical areas and that our dataset contains complementary and previously unexplored mouse thalamic single-nuclei transcriptomes, including mediodorsal thalamus (MD) and intermediodorsal (IMD) thalamus [29].

## RESULTS

### Single-nucleus RNA-sequencing in and around the mouse PVT

To determine the unique transcriptional profiles of individual PVT neurons, single nuclei suspensions were first collected from tissue punches of the PVT and surrounding regions and sequenced at the single-nuclei level (Figure 1a; See Methods) [30]. Next, low-quality nuclei and doublets were removed based on standard criteria (i.e., level of mitochondrial transcripts, number of genes) and integration using the Harmony algorithm was implemented to minimize experimental batch effects (Figure 1—figure supplement 1) [31]. Clustering analyses was performed on 13,220 single-nuclei transcriptomes yielding a total of fourteen clusters visualized by uniform manifold approximation and projection (UMAP) (Figure 1b, c). The nuclei from these clusters were then assigned to seven major cell types based on their differential gene expression of canonical marker genes such as: astrocytes (*Agt*, *A2m*), endothelial (*Flt1*), ependymal (*Tmem212*), microglia (*Cx3cr1*), neurons (*Rbfox3*), oligodendrocyte precursor cells (OPCs) (*Pdgfra*), and oligodendrocytes (*Mal*) (Figure 1d, Supp. Table 1).

**Figure 1.**
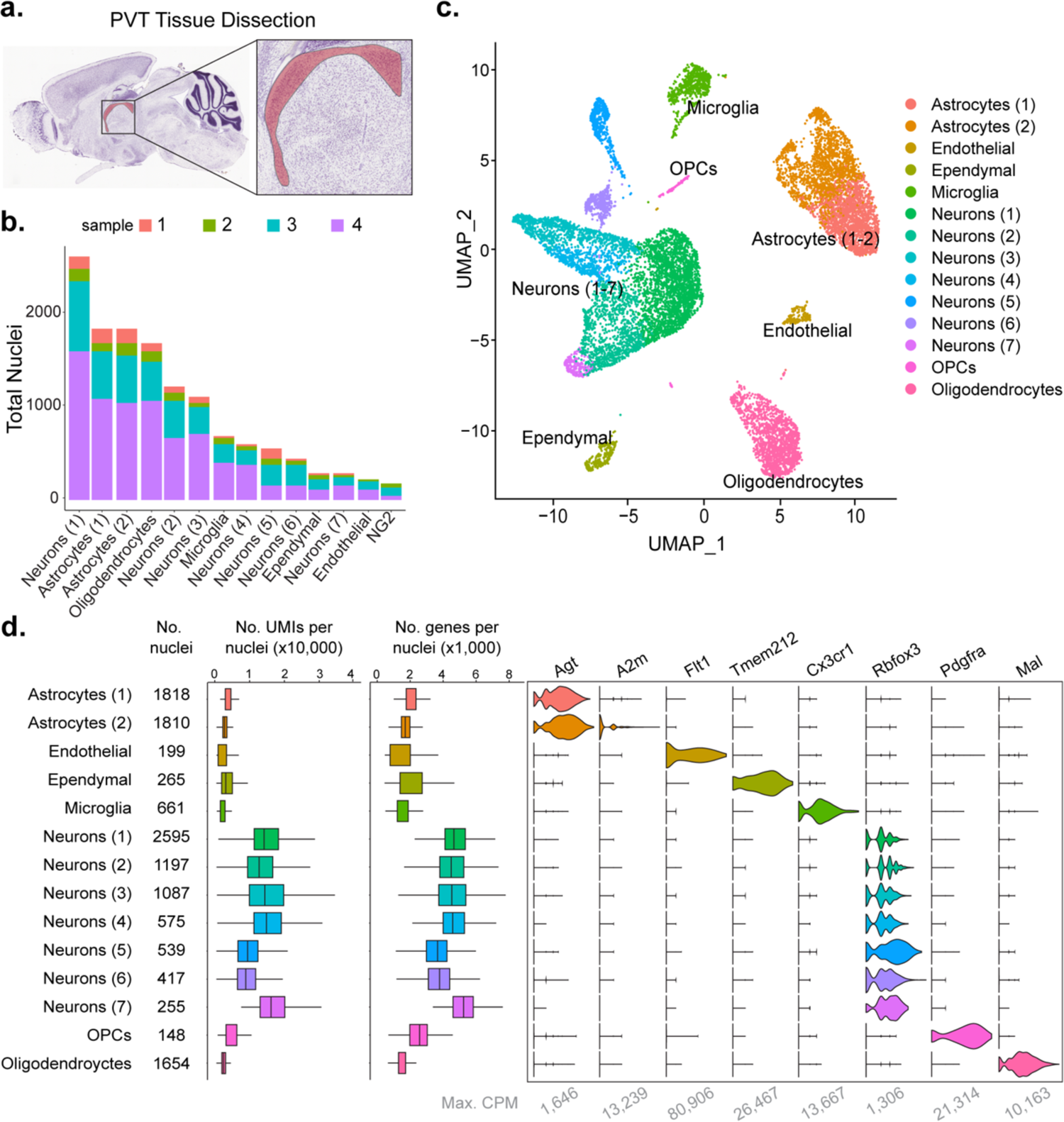
Single-nucleus RNA-sequencing in and around the mouse PVT. **a.** Sagittal view of the PVT (red) illustrating the dissection target location. **b.** Distribution of nuclei from 4 samples across all cell types. **c.** The UMAP plot of all 13,220 nuclei from combined dataset shows 14 cell clusters. **d.** Cell type classification based on expression of marker genes in all 14 clusters. Left: box plot of UMI number in each cell cluster. Middle: box plot of genes detected per cell in each cell clusters. Right: violin plot showing expression profile of marker genes in 14 cell clusters. Max. CPM, maximum counts per million reads. Box plot legend: box is defined by 25th, 75th percentiles, whiskers are determined by 5th and 95th percentiles, and mean is depicted by the square symbol.

To assess neuronal cell types specifically, the seven clusters of neurons (1-7), containing a total of 6,555 nuclei, were then re-clustered for further classification (Figure 1—figure supplement 2a). Clusters were characterized by referencing top gene markers from each cluster with their spatial expression from Allen Brain Atlas ISH data and previous literature (Figure 1—figure supplement 2b, c, mouse.brain-map.org) [32]. From this comparison, we found that PVT neurons (4,067 nuclei; 62.0% of all neurons) expressed markers such as *Gck*, *C1ql3* and *Hcn1*, and were thus represented in neuron clusters 0, 2, 3 and 4 (Figure 1—figure supplement 2c, h) [33–35]. Neurons associated with other brain regions surrounding the PVT and represented in our sample include: MD (cluster 1; 1,003 nuclei, 15.3% of neurons), IMD (cluster 5; 500 nuclei, 7.62% of neurons), principal nucleus of the posterior bed nucleus of the stria terminalis (BSTpr; cluster 6; 496 nuclei, 7.57% of neurons), and habenula (Hb; cluster 7; 489 nuclei, 7.46% of neurons) (Figure 1—figure supplement 2d-g, h, Supp. Table 2). Altogether, we observed eight unique neuronal clusters representing PVT and other adjacent nuclei, almost all of which are thalamic.

### Classification of PVT neuronal subtypes

To explore the cell type heterogeneity of neurons characteristic of the PVT, we removed all non-PVT neurons from subsequent analysis and focused our attention on the characterization of a total of 4,067 PVT nuclei (See Methods). Clustering analysis of PVT neurons revealed 5 neuronal subtypes (PVT1-5), each expressing a unique genetic profile (Figure 2a). A phylogenetic tree was then constructed by generating a distance matrix between clusters in gene expression space. We found that PVT neurons segregated into two major branches (Figure 2b). Specifically, PVT1, PVT2, and PVT5 subtypes were more closely related than PVT3, and PVT4 subtypes. Next, we performed differential gene expression analysis (DGEs) to select top marker genes for each cluster (Figure 2c-h) (See Methods). Since our previous study identified two types of PVT neurons based on their expression of the *Drd2* gene or lack thereof, we compared the expression of *Drd2* across our five molecularly defined subtypes. PVT1 (355 DGEs) and PVT2 (417 DGEs) subtypes expressed *Drd2*, whereas PVT3 (652 DGEs), PVT4 (265 DGEs) and PVT5 (104 DGEs) did not (Figure 2—figure supplement 1). These data suggest that, while as recently reported the PVT can be divided into Type 1 and Type 2 neurons based on *Drd2* expression, this view is oversimplified and instead the PVT contains five molecularly distinct neuronal subtypes [10]. Accordingly, at the molecular level, *Drd2*-positive and *Drd2*-negative neurons can be further divided into two and three subtypes, respectively.

**Figure 2.**
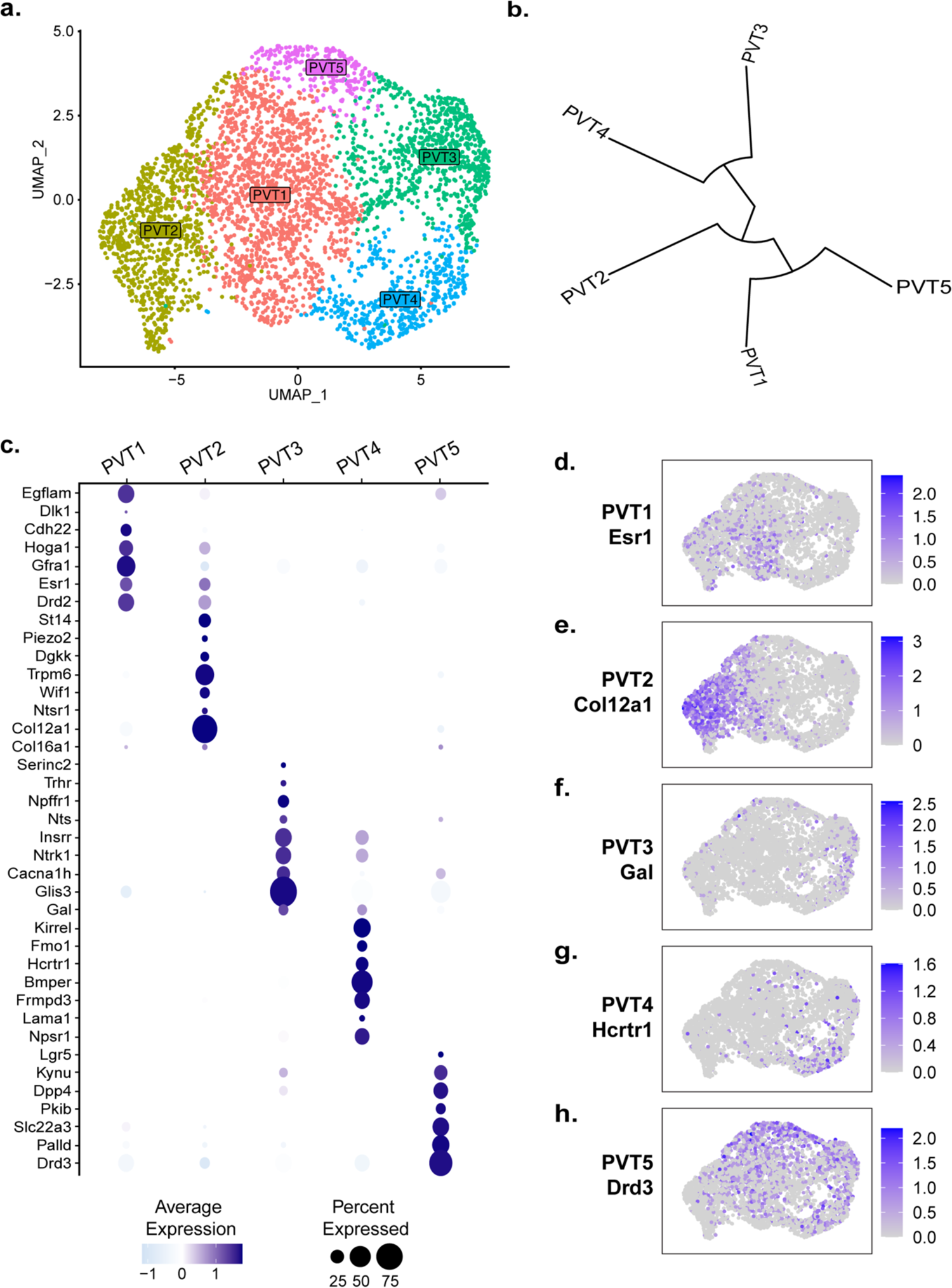
Five transcriptionally distinct neuronal subtypes are found in the PVT. **a.** The UMAP plot of 4,067 PVT neuronal nuclei shows 5 clusters. **b.** Phylogenetic tree depicting cluster relationships based on distance between clusters in gene expression space. **c.** Dot plot of top gene marker average expression across PVT clusters selected based on pct. ratio value. **d-h.** Feature plots of top marker genes (d) Esr1, (e) Col12a1, (f) Gal, (g) Hcrtr1, and (h) Drd3 for each PVT subtype.

### Spatial distribution of 5 PVT neuron subtypes

We next validated the spatial distribution of our five putative PVT neuronal subtypes by employing multiplex ISH assays, where multiple rounds of ISH labeling and cleavage are performed in the same tissue sample. Using this method, we labeled the following cell type-specific markers: *Esr1* (PVT1), *Col12a1* (PVT2), *Gal* (PVT3), *Hcrtr1* (PVT4), and *Drd3* (PVT5) across the antero-posterior axis of the PVT (Figure 3a-g, Supp. Table 3). Following confocal imaging and post-hoc registration, we compared the relative expression of all five marker genes simultaneously across the anterior and posterior PVT. We observed that PVT1/*Esr1* and PVT2/*Col12a1* were significantly expressed in the pPVT, with little to no expression in the aPVT. In contrast, we found that PVT3/*Gal* and PVT4/*Hcrtr1* represented two aPVT biased cell types, which consistent with our previous report [10], are distributed in a gradient like manner with low levels of expression localized to the dorsal pPVT. Finally, PVT5/*Drd3* was expressed across both the aPVT and pPVT in a pan-PVT like manner (PVT5^pan^) (Figure 3i-k). We also performed quantitative analyses of our multiplex ISH labeled samples to determine the distribution and number of positive cells for each transcript and generated a normalized transcript expression matrix (Figure 3h). From this matrix, we computed the Euclidian distances between gene markers and performed hierarchical clustering of our data. Consistent with our sequencing results, we found that PVT3/*Gal* and PVT4/*Hcrtr1* expressing cells are more closely related than PVT1/*Esr1*, PVT2/*Col12a1*, and PVT5/*Drd3* expressing cells (Figure 3—figure supplement 1a). Notably, hierarchical clustering based on our multiplex ISH data suggested that PVT1 and PVT2 are more closely related to each other than to PVT5. This difference from the sequencing phylogenetic tree is likely a result of how the distance matrices between the spatial data and the sequencing data are calculated. The spatial expression matrix compares one gene marker per cluster, providing an overview of the spatial relationship of each cluster, whereas the sequencing phylogenetic tree is generated using gene expression across all cells in a cluster, yielding a higher resolution view of each cluster’s transcriptional relationship [36]. Together, these provide information on both the spatial context and transcriptional relationship of PVT cell types [37].

**Figure 3.**
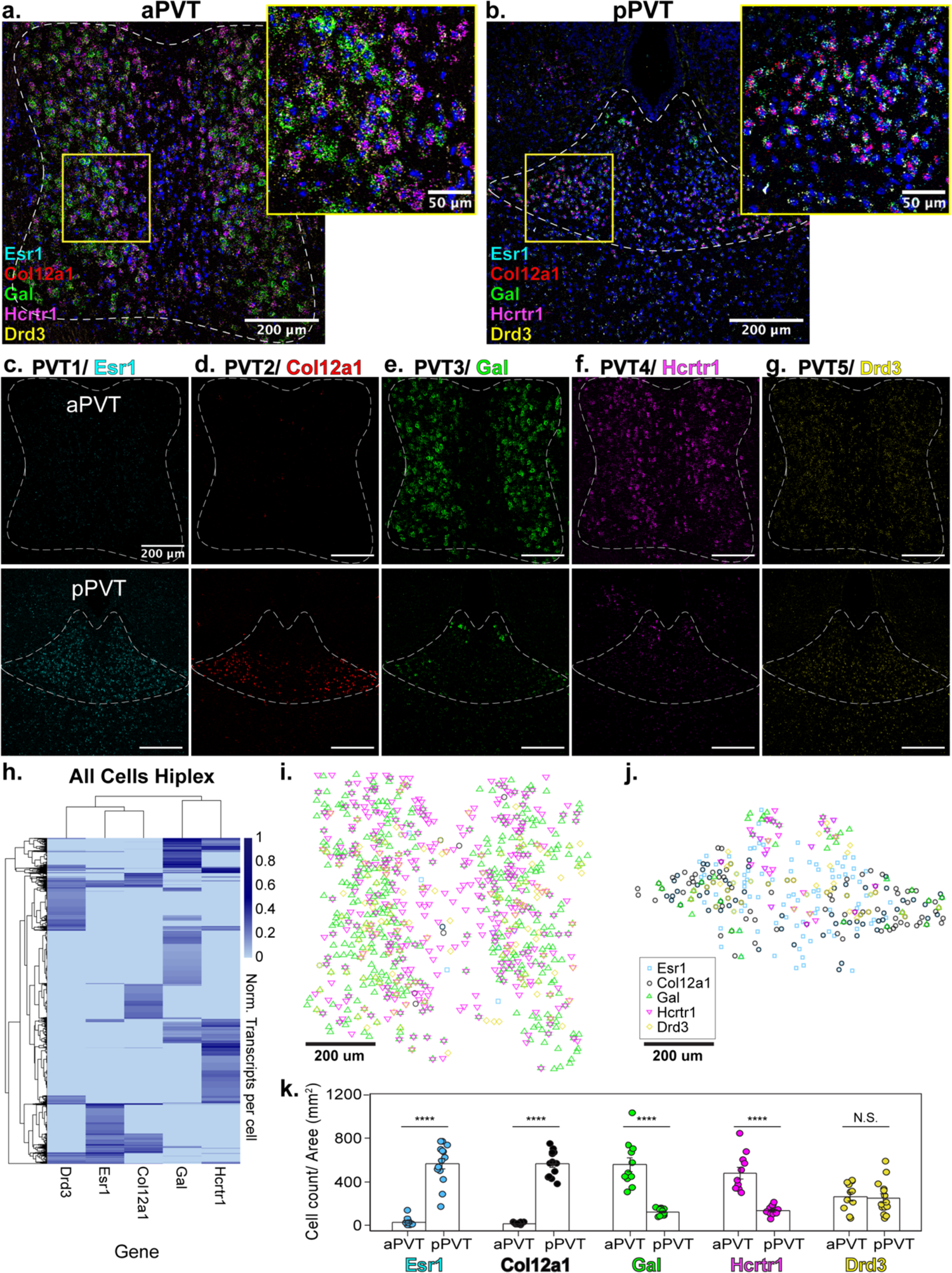
Mapping of 5 PVT subtypes reveals segregated spatial gradients. **a.** RNA in situ hybridization in anterior PVT (white dotted outline) with closeup insert (right, yellow square) labeling one gene marker from each subtype: *Esr1* (light blue), *Col12a1* (red), *Gal* (green), *Hcrtr1* (magenta), and *Drd3* (yellow). **b.** RNA in situ hybridization in posterior PVT (white dotted outline) with closeup insert (right, yellow square) labeling one gene marker from each subtype: *Esr1* (blue), *Col12a1* (red), *Gal* (green), *Hcrtr1* (magenta), and *Drd3* (yellow). **c-g.** RNA in situ hybridization showing aPVT (top) and pPVT (bottom) of (c) *Esr1* from PVT1 subtype, (d) *Col12a1* from PVT2 subtype, (e) *Gal* from PVT3 subtype, (f) *Hcrtr1* from PVT4 subtype, (g) *Drd3* from PVT5 subtype. DAPI (blue). Scale bar 200um; all images are 20x representative confocal images with brightness and contrast adjusted depicting expression patterns found in all sections from N=3 animals. **h.** Heatmap of gene expression matrix with rows showing normalized transcripts per cell and columns showing gene markers from each PVT subtype; Left dendrogram displays hierarchical clustering by cell, top dendrogram displays hierarchical clustering by gene marker, right bar shows heatmap legend. **i-j.** Coordinates of positive cells from aPVT (i) and pPVT (j) with *Esr1* (blue square), *Col12a1* (black circle), *Gal* (green triangle), *Hcrtr1* (magenta downward triangle), *Drd3* (yellow circle) shown in legend. **k.** Bar graphs of number of positive cells over the area (mm^2^) per section of *Esr1* (blue; aPVT: 30.71 ± 11.73; pPVT: 179.78 ± 48.05; \*\*\*\**P*=0.0000000015, two-sided Paired sample t-test), *Col12a1* (black; aPVT: 15.85 ± 2.95; pPVT: 563.13 ± 30.37; \*\*\*\**P*=0.000000000000071, two-sided Paired sample t-test), *Gal* (green; aPVT: 557.26 ± 63.47; pPVT: 123.19 ± 8.72; \*\*\*\**P*=0.000000092), *Hcrtr1* (magenta; aPVT: 478.53 ± 52.05; pPVT: 133.62 ± 10.76; \*\*\*\**P*=0.00000021), and *Drd3* (yellow; aPVT: 265.97 ± 38.52; pPVT: 250.43 ± 41.18; N.S., not significant, P=0.79) in aPVT and pPVT. Data from N=3 animals shown as mean ± S.E.M. Each datapoint represents one section.

Plotting the locations of *Esr1*, *Col12a1*, *Gal*, *Hcrtr1*, and *Drd3* positive cells reveal a unique spatial segregation within the anterior-biased and posterior-biased clusters (Figure 3i-j). PVT4/*Hcrtr1* cells were localized more antero-medially (PVT4^AM^), whereas PVT3/*Gal* cells were expressed more antero-laterally (PVT3^AL^). Also, while PVT1/*Esr1* cells were located across the dorso-ventral axis of the pPVT (PVT1^P^), PVT2/*Col12a1* cells were restricted to the ventral portion of the pPVT and seemed to have strongest expression at the bottom “edge” of the pPVT (PVT2^edge^). To better visualize the spatial distribution among gene markers, we separately plotted pan-PVT (PVT5^pan^) cells with either the pPVT (PVT1^P^, PVT2^edge^) or the aPVT (PVT3^AL^, PVT4^AM^) biased clusters, and examined the percentage of overlap between positive cells amongst PVT subtypes (Figure 3—figure supplement 1b-g). Altogether, we find that, while there are varying degrees of overlap at the single cell level, these gene markers largely represent spatially discrete cell types.

Importantly, while there are several markers that appear exclusive to PVT2^edge^, some of the markers for PVT1^P^ were also expressed to a weaker degree in PVT2^edge^ (Figure 2c). Thus, to further confirm the nature of spatial distribution of our five PVT subtypes, we selected a second set of markers from either neuromodulator receptor or ion channel genes for each of the aPVT biased (PVT3^AL^, PVT4^AM^) and pPVT (PVT1^P^, PVT2^edge^) biased subtypes and compared their spatial distribution to those of our prototypical PVT cluster markers (*Esr1*, *Col12a1*, *Gal*, and *Hcrtr1*) using multiplex ISH (Supp. Table 3). For this, we selected *Drd2*, *Chrm2*, *Npffr1*, and *Npsr1*, as representative markers for subtypes PVT1^P^, PVT2^edge^, PVT3^AL^, and PVT4^AM^, respectively. Labeling of this second set of markers across the anterior and posterior PVT revealed a similar spatial distribution as our initial selected markers (i.e., AM, AL, P, edge) with only 30% or less overlap at the single-cell level among PVT subtype marker sets (Figure 3—figure supplement 2, 3). These findings indicate that there is a high amount of diversity in gene expression even within the same PVT subtype and may suggest that, while there are only five molecular subtypes, cells within the same PVT subtype could have non-overlapping functions tied to differential gene expression.

Overall, in addition to validating the results of the snRNA-seq data, spatial mapping of our five PVT subtypes demonstrates the existence of spatially segregated, and molecularly distinct neuronal subpopulations of the PVT. As such, our data highlight a new feature of PVT cell type organization in which the PVT is structured by a combination of gradients: antero-posterior, dorsal-ventral, and medio-lateral. This arrangement of molecularly distinct cellular subtypes, wherein genetic identity and spatial distribution go hand in hand is a recurrent feature of the mammalian brain [29, 38–42]. Importantly, molecular gradients in other thalamic nuclei, namely the thalamic reticular nucleus (TRN) and MD, have been linked to functional differences [43–45]. The existence of a similarly close relationship between molecular identity and function among neurons of the PVT highlights how the discovery of additional molecular gradients may provide a framework for redefining the functional organization of the PVT [10, 46].

### The 5 PVT subtypes have unique profiles of expression for neuromodulator receptors, neuromodulators, and ion channels

The PVT is a site of convergence of dense peptidergic innervation from cortical, hypothalamic, and hindbrain areas [4, 20, 47–53]. As such, previous studies have indicated a segregated distribution and differential effects of neuromodulatory innervation and receptors along the antero-posterior axis of the PVT [54–59]. To examine differences in genes that encode for proteins that regulate cellular excitability or function, we compared the expression of top differentially expressed neuromodulator receptors, neuromodulators, and ion channels across our five PVT subtypes (Figure 4a-c, Supp. Table 3). In addition to many neuromodulatory systems previously reported in the PVT, such as the orexin/hypocretin system (*Hcrtr1*/*Hcrtr2*) [48], dopamine system (*Drd2*) [60, 61], and endocannabinoid system (*Cnr1*) [62, 63], our data revealed the existence of many previously uncharacterized neuromodulatory receptors and ion channel subunits across the five PVT subtypes. For example, *Npffr1,* the gene encoding neuropeptide FF receptor 1, is represented in the PVT3^AL^ subtype, whereas *Chrm2,* the gene encoding for muscarinic acetylcholine receptor M2, is expressed in the PVT2^edge^ subtype (Figure 4a). Furthermore, within the same neuromodulatory system, we found differences in receptor family expression across PVT types. For instance, dopamine receptor genes *Drd1* and *Drd3* were identified in PVT3 ^AL^ and PVT5 ^pan^ subtypes, respectively, whereas *Drd2* was found in PVT1^P^ and PVT2 ^edge^ subtypes. Similarly, glycine receptor genes *Glra1*, *Glra2*, *Glra3* were differentially represented across PVT2, PVT3, and PVT4 subtypes, respectively (Figure 4a).

**Figure 4.**
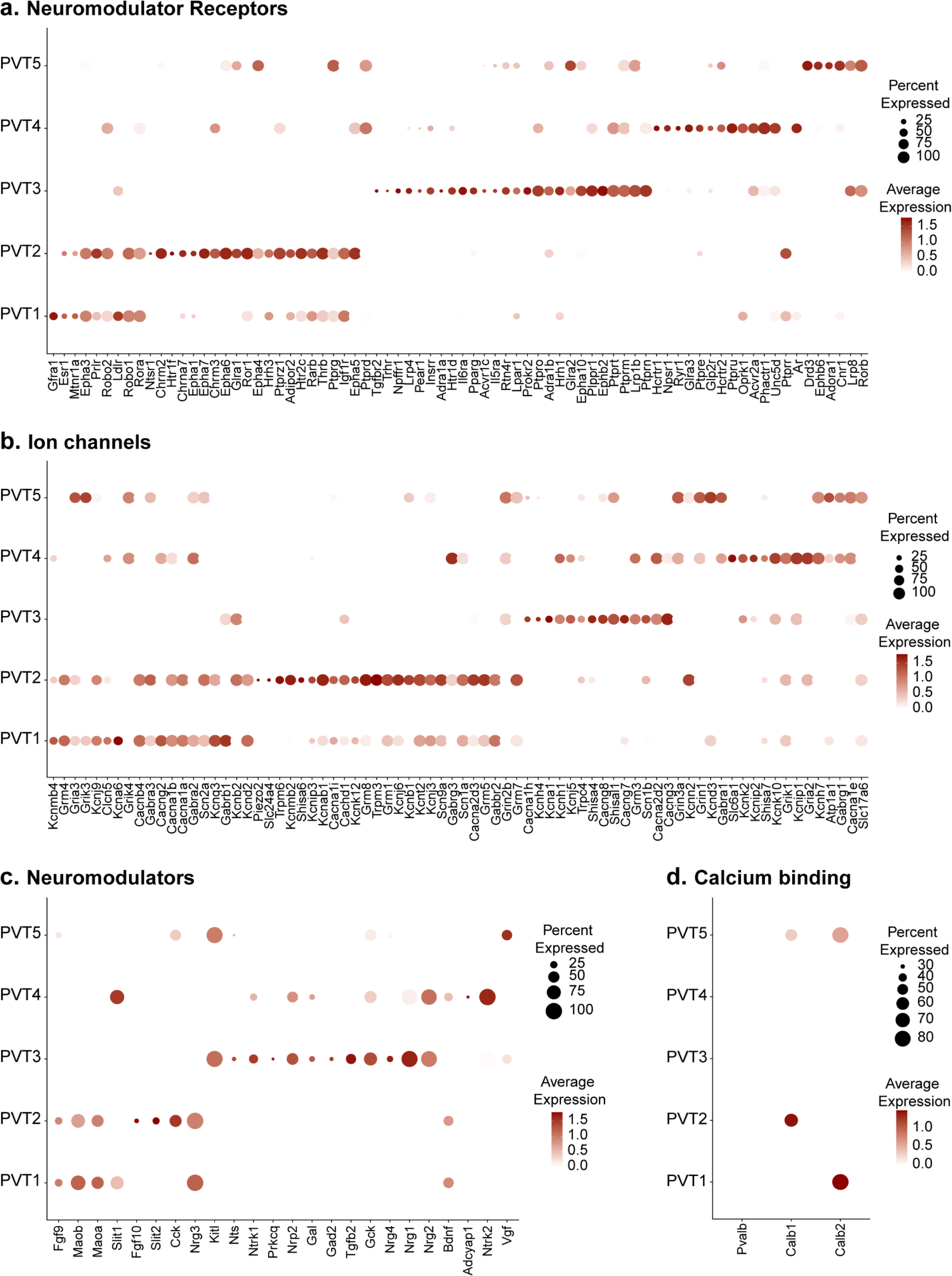
Five PVT subtypes have diverse neuromodulator receptor, neuromodulator, and ion channel expression. **a.** Dot plot depicting neuromodulator receptor gene expression across five PVT subtypes. **b.** Dot plot depicting ion channel gene expression in across five PVT subtypes. **c.** Dot plot depicting neuromodulator gene expression across five PVT subtypes.**d.** Dot plot depicting calcium binding gene (column) expression across five PVT subtypes. Legend: top, percent of nuclei in each cluster expressing a given gene; bottom, color intensity corresponding to average gene expression level.

Given that differential expression of ion channel genes may be linked to divergence in the intrinsic membrane properties of PVT neurons, we compared ion channel gene expression across PVT types and observed substantial heterogeneity (Figure 4b) [35, 64–66]. T-type calcium (Ca^2+^) channels have been documented to contribute to the unique membrane properties and overall excitability of PVT neurons [35, 64, 65, 67]. Interestingly, we found that while T-type Ca^2+^ channel subunit gene *Cacana1h* was most robustly expressed in PVT3^AL^ subtype, *Cacna1i* was differentially expressed in PVT1^P^ and PVT2^edge^ subtypes (Figure 4b). Similarly, we observed that voltage-gated potassium channel genes *Kcna1* and *Kcnb1* were differentially expressed in PVT3^AL^ and PVT2^edge^ subtypes. Such differential expression of voltage-gated potassium channels may further underlie variability in the membrane properties of PVT neurons (Figure 4b) [35]. Altogether, these findings highlight how our transcriptomic database may serve as a resource to identify key markers for future functional investigations, including genetic targeting of specific cell types.

Finally, we compared the gene expression of certain Ca^2+^ binding proteins, including *Pvalb*, *Calb1*, and *Calb2* across the five PVT subtypes (Figure 4d). *Calb2*, the gene encoding calretinin, is a common gene marker used to identify PVT neurons that have recently been implicated in arousal signaling [8, 9]. Here, we find that *Calb2* expression is found in PVT1^P^ and PVT5^pan^, which is consistent with previous reports that expression of this gene is found across the PVT [8, 9]. *Calb1*, the gene encoding calbindin 1 (CB), and *Pvalb*, the gene encoding parvalbumin (PV), are classical markers used in a thalamic classification system to segregate between what are known as ‘core’ and ‘matrix’ thalamic relay cells [68]. PV-expressing ‘core’ thalamic cells are defined by their topographical and dense projections to the middle layers of defined cortical regions, whereas CB-expressing ‘matrix’ thalamic cells instead send diffuse projections across unrestricted superficial cortical fields [69–71]. From these projection patterns, ‘core’-containing thalamic regions have often been deemed first order thalamic nuclei, such as principal sensory thalamic relay nuclei [72]. On the other hand, ‘matrix’-containing regions that lack ‘core’ cells are often referred to as higher-order thalamic nuclei, such as the midline and intralaminar nuclei [72]. Consistent with descriptions of the PVT as a higher-order midline thalamic structure, all five PVT subtypes lacked *Pvalb* expression (Figure 4d) [68, 71]. Notably, *Calb1* expression was only found in PVT2^edge^ and PVT5^pan^ subtypes, with higher expression in PVT2^edge^. This result alongside other studies lends support to the notion that the core and matrix dichotomy is not a sufficient classification system to encompass all thalamic cell types [29, 73, 74].

### Cross-validation with Thalamoseq dataset reveals overlap between PVT subtypes and thalamic cortical projectors

To cross-reference our snRNA-seq dataset of the mouse midline thalamus, we performed integration using RPCA together with the recently published ThalamoSeq RNA-Seq atlas (thalamoseq.janelia.org), which contains single-cell sequenced thalamic cells that project to either motor, auditory, visual, somatosensory, or prefrontal cortices (PFC) (Figure 5, Figure 5—figure supplement 1) [29, 36]. Upon clustering analysis and plotting the UMAP of the integrated dataset with all midline thalamic and BSTpr neurons, we found that our midline thalamic dataset and the ThalamoSeq dataset contained mostly non-overlapping transcriptomes (Figure 5—figure supplement 1).

**Figure 5.**
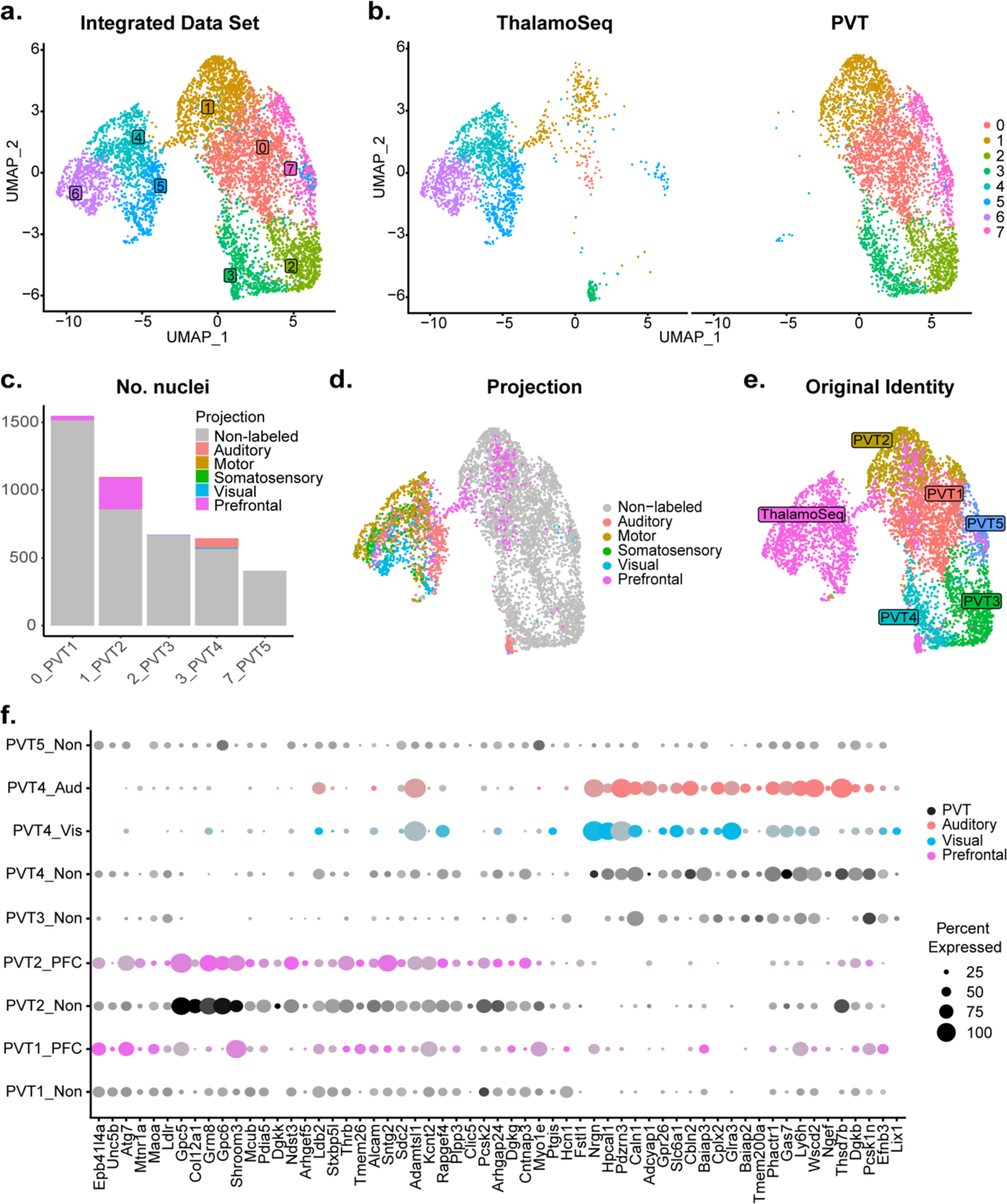
Cross-validation with Thalamoseq dataset reveals overlap between PVT subtypes and thalamic cortical projectors. **a.** The UMAP plot of 6,0397 cells or nuclei from combined datasets annotated by cluster identity. **b.** The same cells/nuclei separated by study of origin: Thalamoseq (left) and present study (right) **c.** Proportion of nuclei from each cortical projection target in the clusters that represent the molecular PVT subtypes from the integrated dataset: 0=PVT1, 1=PVT2, 2=PVT3, 3=PVT4, 7=PVT5. **d.** The same cells/nuclei colored by cortical projection target. **e.** The same cells/nuclei colored by original identity. **f.** Dot plot of PVT representing clusters (0,1,2,3,7) in the integrated dataset depicting expression of genes obtained from the conserved marker analysis (Supp. Table 5) split by projection identity.

Next, we performed integration of the ThalamoSeq dataset and PVT neurons specifically (Figure 5a-b). By mapping the integrated dataset based on cortical projection location from the ThalamoSeq cells or their original cluster identity (either PVT1-5 or ThalamoSeq), we could observe the cluster relationship between ThalamoSeq cortical projection location and our five PVT molecular subtypes (Figure 5c-e). DGE analysis on the integrated dataset revealed that our original PVT subtypes (PVT1-5) could be mapped onto clusters 0, 1, 2, 3, and 7, respectively, of the integrated dataset (Supp. Table 4, Figure 5a, c). Following this, comparing the proportion of cortical-projectors in the integrated dataset showed that the clusters representing PVT1^P^ and PVT2^edge^ contained overlap with PFC-projectors, whereas, unexpectedly, the cluster representing PVT4^AM^ had a small amount of overlap with auditory and visual projectors (Figure 5b). The clusters representing PVT3^AL^ and PVT5^pan^ exhibited little to no overlap with any cortical projectors (Figure 5b). Notably, the greatest amount of overlap was observed between ThalamoSeq PFC-projectors and the PVT2^edge^ subtype, represented in cluster 1 of the integrated dataset. Next, we performed conserved marker analysis across these five integrated clusters and confirmed that *Col12a1* is a conserved marker across PFC-projectors and PVT2^edge^ cells (Figure 5f, Supp. Table 5). These findings suggest that the *Col12a1*-positive PVT2^edge^ subtype represents a subpopulation of neurons that project to the PFC, which is consistent with prior studies indicating that pPVT neurons are anatomically linked to the prelimbic cortex (PL) in a cortico-thalamocortical loop [10, 17, 19, 75]. Accordingly, anatomical tracing studies from the Mouse Connectome Project revealed that PVT efferent and afferent connectivity with the PL is restricted to the ventral edge of the pPVT (Figure 5—figure supplement 2, www.mouseconnectome.org) [76]. Given that the cluster representing PVT4^AM^ overlapped with auditory and visual cortical projectors, we next employed the Allen Brain Mouse Connectivity Atlas Target Search Tool to look for auditory and visual areas that receive innervation by the PVT (Figure 5—figure supplement 3, connectivity.brain-map.org). Consistent with the results from our comparative analysis with the ThalamoSeq dataset, this search revealed four experiments demonstrating that anterior-biased thalamic regions including the aPVT (but not the pPVT) send projections to known auditory and visual cortical areas, such as the temporal association cortex (Figure 5—figure supplement 3). Collectively, the integrative analysis performed here supports that different PVT subtypes, namely PVT1^P^, PVT2^edge^, and PVT4^AM^, share conserved markers with different thalamo-cortical projectors.

## DISCUSSION

Despite being critical nodes in linking subcortical and cortical networks, to date, models of the spatial and molecular organization of midline and intralaminar thalamic nuclei have remained scarce [10, 19, 20, 46, 77]. Recently, our group and others have demonstrated that, contrary to the traditional discrete parcellations of thalamic nuclei, functionally distinct thalamic domains are organized in continuous neuronal gradients. Specifically, the PVT, TRN, and MD thalamus have each been shown to contain at least two genetically defined cell types that are organized in opposing spatial gradients and reflect distinct anatomical connectivity, electrophysiological signatures, and functional contributions [10, 43–45]. To expand this framework, in the present study, we generated an in-depth single-nuclei resolution transcriptomic atlas of the PVT that revealed the existence and spatial distribution of five previously undescribed neuronal subpopulations. Importantly, these PVT subtypes are organized in a combination of topographical gradients (e.g., medio-lateral, antero-posterior, dorso-ventral), which, to our knowledge, has not yet been reported in another midline thalamic area. However, given that thalamic pathways consist of repeated molecular architectures [29], it is likely that this feature of multiple gradients is conserved and may serve as a basis for new conceptual models across midline and intralaminar thalamic organization.

From our molecular and spatial mapping, we found that the five subtypes can be split into two major branches: (1) PVT1^P^, PVT2^edge^, PVT5^pan^ and (2) PVT3^AL^, PVT4^AM^. These two branches consist of the posterior-biased and pan-PVT, and anterior-biased subtypes, respectively, indicating that one major axis of separation between these two branches lies in their rostral-caudal distribution. While this notion is in partial agreement with classical anatomical studies delineating anterior and posterior PVT subregions, our study supports the ongoing revision of this terminology and fills a critical gap in our knowledge of the genetic heterogeneity of subpopulations within the PVT [7].

From classical tracing studies, the pPVT selectively innervates PL, nucleus accumbens core and ventromedial shell, central and basolateral amygdala, and lateral BST [17, 19]. In line with this distinct connectivity, functional correlates implicate the pPVT as a key node in networks of aversive and reward motivated behaviors, including stress and fear processing, and food- and drug-seeking. Importantly, our observation that there are two gradients of posterior-biased subpopulations—PVT1^P^ and PVT2^edge^— gives rise to the question of whether these subtypes contribute to discrete functions. Of note, our integrative analysis with the ThalamoSeq atlas showed that PVT2^edge^ cells in particular belong to a group of *Col12a1*-expressing PFC-projectors [29]. This result is consistent with classical tracing studies placing the pPVT in a reciprocal network with the PFC, and more specifically the PL [17, 19, 75]. Intriguingly, anterograde viral tracing from multiple studies reveals that prelimbic afferents to the pPVT are spatially distributed in a manner that resembles the ventroposterior location of PVT2^edge^ cells, lending support to the idea that PVT2^edge^ cells may be preferentially integrated into a cortico-thalamocortical pathway that mediates its role in cognitive processes related to emotional and motivational states (Figure 5—figure supplement 2) [76, 78, 79]. Indeed, prefrontal projections to the PVT guide the expression of emotional memory-associated behaviors such as conditioned fear and cue-reward food seeking [78–83]. In this context, the PVT2^edge^ cells may participate in the integration of cortical influence onto downstream subcortical targets to bias behavioral responding during learning and goal-oriented tasks where top-down modulation is necessary [78–83]. In contrast, PVT1^P^ neurons integrated to a lesser extent with PFC-projectors (Figure 5c). Interestingly, the medial portion of the pPVT, which lacks PVT2^edge^ expression and is mostly populated by PVT1^P^ neurons, is heavily innervated by subcortical inputs which may participate in biasing behavior toward stereotyped emotional reactions like those observed amid imminent environmental threats [84–88]. Additionally, PVT1^P^ neurons specifically express *Esr1* (estrogen receptor 1), a cell marker for ventromedial hypothalamic (VMH) neurons that control aggression and mating, which are stereotyped and conserved behaviors under the control of subcortical networks [89–92]. Whether *Esr1* neurons in the PVT contribute to the control of aggressive and mating behavior is not known; however, the PVT receives heavy afferents from *Esr1*-expressing VMH neurons, anatomically linking these two structures and prompting further investigation [93]. Altogether, we propose that PVT2^edge^ may consist of a genetic subpopulation that is preferentially involved in aspects of behavior requiring cortical processing, whereas the PVT1^P^ subtype may contain neurons differentially recruited by tasks involving subcortical or cortical networks depending on their location.

In addition to two posterior biased subpopulations, we discovered the existence of two anterior biased molecularly distinct gradients: antero-lateral PVT3^AL^ and antero-medial PVT4^AM^. Previous tracing studies indicate that the aPVT preferentially innervates the dorsomedial shell of the NAc and suprachiasmatic nucleus of the hypothalamus [17–19], and functional studies have linked aPVT to arousal, cardioprotection, and the inhibition of reward-seeking behaviors [10, 28, 55, 94, 95]. Notably, reports of aPVT contributions to arousal signaling are complex, whereby activation of aPVT neurons has been shown to drive both increases and decreases in wakefulness [8, 10]. Specifically, *Gal* (galanin)-expressing neurons were proposed to belong to a subclass of aPVT neurons that antagonizes arousal and increases NREM sleep [10], which contrasts findings showing that *Calb2*-expressing aPVT neurons instead promote arousal and increase wakefulness [8, 9]. These differences might find explanation in the present study where we show that *Gal* is a cell marker of PVT3^AL^ subtype, whereas *Calb2* is expressed in both PVT1^P^ and PVT5^pan^ subtypes, with higher expression levels in PVT1^P^, suggesting that it is spatially distributed across the entire PVT with a bias toward the pPVT (Figure 4b). As such, it seems plausible that the PVT exerts bidirectional control over arousal and wakefulness through molecularly and spatially diverse subpopulations [8–10]. Interestingly, *Gal*-expressing PVT neurons were reported to preferentially target the infralimbic cortex (IL) and are likely reciprocally connected with IL [10] (Figure 5—figure supplement 4). This result suggests that, like the posterior biased PVT2^edge^ subtype, PVT3^AL^ neurons may be preferentially integrated in a cortico-thalamocortical pathway that mediates cortical influence over emotional and motivated processes. In support of this, a recent study showed that manipulation of aPVT-projecting IL neurons could alter levels of physiological arousal in response to tail shock in mice [10].

It is important to highlight that, contrary to findings related to PVT2^edge^ neurons and PVT1^P^, RPCA-based integration with the ThalamoSeq dataset did not reveal PFC-projectors among other PVT subtypes. One potential explanation for the lack of overlap between other PVT clusters and the ThalamoSeq atlas could lie in the retrograde labeling and sample collection method used [96]. Indeed, the authors of ThalamoSeq reported that their single cell collection was biased against midline thalamic nuclei due to issues related to sparse retrogradely labeled cells and thus, lack of overlap should not preclude other PVT subtypes, including PVT3^AL^, as PFC-projectors [29].

We also report here that the glucokinase (*Gck*) gene is most highly expressed in the PVT3^AL^ subtype, potentially linking recent reports of the unique attributes of glucokinase-expressing aPVT neurons (aPVT^Gck^) to the PVT3 subtype [27, 33]. NAc-projecting aPVT^Gck^ neurons were shown to be activated by hyperglycemic conditions and negatively modulate sucrose-seeking behavior [27]. This result is consistent with other reports showing that inactivation of aPVT projections to the NAc leads to increased food-seeking behavior, particularly when reward is unavailable, suggesting that activity in this pathway controls behavioral suppression [28, 95, 97]. Importantly, in contrast to the aPVT-NAc pathway, some studies demonstrate that pPVT projections to the NAc promote food seeking behavior [14, 15]. Similar to the PVT’s bidirectional control over arousal, we propose that these opposing contributions to reward-seeking originally attributed to binary subregions may instead be a feature of molecularly distinct PVT gradients biased toward either the aPVT or pPVT. Notably, there is some *Gck* expression found in both PVT4^AM^ and PVT5^pan^, possibly implicating these two subtypes in the inhibition of food-seeking behavior as well. Collectively, we propose that PVT3^AL^ neurons are likely involved in the negative modulation of arousal and reward-seeking behavior, thereby enabling the PVT to engage in flexible control of emotional and motivated processes.

Based on the top markers derived from the current study, PVT4^AM^ neurons appear to represent a lesser studied PVT subtype. Although these neurons highly express the orexin receptor 1 gene (*Hcrtr1*), and orexinergic influence in the PVT has been linked to anxiety, stress, arousal, and food- and drug-related behaviors [15, 98–103], evidence linking orexin receptor 1 (Ox1R) in the PVT to these processes is scarce [103–107]. Instead, most research reports on the role of orexin receptor 2 (Ox2R), encoded by the *Hcrtr2* gene, whose expression is found in both PVT4^AM^ and PVT5^pan^ subtypes (Figure 4a). Specifically, activation of Ox2Rs in the pPVT has been documented to increase anxiety behaviors, promote arousal, drive food-seeking, and facilitate pro-addictive drug-seeking behavior [15, 55, 59, 98, 102, 108, 109]. While studies specifically interrogating the function of PVT Ox1Rs are limited, shRNA knockdown of PVT Ox1Rs decreased hedonic feeding behavior in sated rats but had no effects on behavioral responding in a progressive ratio task—an instrumental task that measures both motivation and learning [103, 110]. The authors of this study suggested that Ox1Rs in the PVT support hedonic feeding, but do not affect the motivation for reward-seeking in instrumental tasks, potentially implicating Ox1Rs in food-consummatory behavior but not in food-seeking [103]. This is supported by a separate study showing that systemic blockade of Ox1Rs similarly decreased cue-induced feeding in sated rats, a result that was intriguingly correlated with an increase in cFos expression in the PVT [104]. In contrast, a recent study showed that blockade of Ox2R, but not Ox1Rs, in the pPVT decreased lever-pressing in an instrumental cue-driven reward seeking task in hungry rats [15]. Thus, it is possible that PVT Ox1R and OxR2 (and as such different PVT sectors) may be differentially recruited based on the motivational demand of the task at hand (e.g., food consumption vs. food seeking) [15, 111]. One surprising observation generated by the RPCA-based integration of our dataset with the ThalamoSeq atlas, is the notion that some neurons within PVT4^AM^ may send anatomical projections to auditory and visual cortical areas (Figure 5c, f). This assertion is further supported by anterograde tracing studies showing that the aPVT projects to auditory and visual areas that have been implicated in or suggested to regulate limbic functions (Figure 5—figure supplement 4) [112–114]. Given the PVT’s role in emotional salience processing, these findings raise the possibility that the PVT4^AM^ participates in modulating emotional responses to salient stimuli in part through its interaction with limbic cortical areas beyond the PFC [10, 115–117].

The PVT5^pan^/*Drd3* subtype was revealed following the removal of non-PVT neurons and was the smallest subpopulation of our five subtypes, with shared expression of several neuroactive and ion channel genes as the other clusters (Figure 4). Even so, we found that this cluster demonstrates a transcriptionally distinct profile and expresses *Drd3* (dopamine receptor 3; D3R) (Figure 2c, Figure 3g). Interestingly, *Drd1* (dopamine receptor 1) is highly differentially expressed in the PVT3^AL^ subtype and *Drd2* is found in PVT1^P^/PVT2^edge^, as previously mentioned (Figure 4a). This raises the question of whether the dopaminergic system may exert differential effects over PVT circuits through its molecularly distinct subtypes. In support of this notion, D2R-dependent dopaminergic transmission in the pPVT was associated with increased sensitivity to stress [61], whereas D3R activity in the PVT has been suggested to mediate the action of psychostimulants [118]. However, whether these reported functions are exclusive to the D2R and D3R in the PVT has not been examined given that some agonists, such as quinpirole, may bind both D2Rs and D3Rs with different affinities [119]. For example, infusion of the D2-like agonist quinpirole in the PVT of mice precipitates arousal from anesthesia, but whether quinpirole actions on D2R or D3R (or both) mediate these effects remains unclear [119–121]. Nonetheless, an important spatial feature of genes found within the PVT5 subtype is that they appear to be expressed across the entire axis of the PVT. Whether this corresponds to a uniform function across the PVT is not known. However, in support of this, we find that PVT5 neurons also selectively express *Cnr1* (cannabinoid receptor 1; CBR1), which is found across the PVT (Figure 4a) [122]. Reports suggest that CBR1s modulate PVT neuronal excitability during low threshold Ca^2+^ spiking (LTS), which, in part, are thought to contribute to burst firing and synchronized oscillatory activity underlying sleep-wake behaviors [62, 63]. Notably, LTS is a general feature found across PVT neurons, which may indicate that the PVT5 subtype contains neurons that are functionally related [64, 65].

Overall, we report large variations in neuromodulator system and ion channel gene expression across PVT subtypes (Figure 4c), There is a formative body of evidence supporting that PVT neurons express unique intrinsic properties that undergo diurnal fluctuations associated with changes in gene expression [35]. PVT neuronal firing properties are also documented to be affected by a variety of neuromodulators, which may contribute to differences in their electrophysiological signatures [35, 64]. Additionally, neuromodulator influence in the PVT is higher than other neighboring thalamic regions, perhaps owing to its selective innervation by the hypothalamus and brainstem [20, 47, 123–125]. Given that each subtype contains differential expression of neuromodulator receptor systems and ion channels, the previously reported effects of neuromodulators on the intrinsic properties of PVT neurons are likely to be diverse across subtypes [56, 64, 66, 126–131]. To examine how PVT neurons undergo experience-dependent changes in gene expression, future investigations should include high throughput RNA-sequencing of the PVT following different biological conditions with the goal of comparing them with our atlas to identify potential biomarkers for therapeutic targeting. Indeed, based on the role of the rodent PVT in emotional processing and motivated behaviors, there is growing interest in the human PVT as a putative target for affective and substance abuse disorders, particularly since the human PVT has similar connectivity with the rodent PVT [132]. Excitingly, the existence of some markers (e.g. *Drd2*, *Drd3*) reported in our present atlas have already been corroborated in the human PVT, supporting that our atlas may provide conserved gene markers across species [133].

In conclusion, while there have been some investigations into the molecular organization of the PVT, here we have provided the first comprehensive molecular and spatial profiling of cell types across the entirety of the PVT [29, 33, 134]. The PVT is a critical integrative node linking circuits of internal and external processing with those involved in arousal signaling, as well as instrumental and Pavlovian behavioral control [2–4, 135]. Our discovery of five spatially segregated PVT neuronal subtypes should serve as a basis for resolving previously conflicting observations and offer a starting point into new mechanistic inquiries as to how PVT offers control over its diverse range of functional correlates.

## METHODS

### Mice

All procedures were performed in accordance with the *Guide for the Care and Use of Laboratory Animals* and were approved by the National Institute of Mental Health (NIMH) Animal Care and Use Committee. Mice used in this study were housed under a 12-h light-dark cycle (6 a.m. to 6 p.m. light), with food and water available *ad libitum*. The following mouse lines were used: C57BL/6J (The Jackson Laboratory). Male mice 8-12 weeks of age were used for all experiments.

### Single-nuclei RNA-Sequencing

#### Sample Collection and Single nucleus RNA sequencing

Tissue punches from the entire antero-posterior PVT from four P90 adult male C57BL/6J mice (12 weeks) were dissected and pooled together for each individual sample and a total of four samples were collected. Nuclei were obtained using the mechanical-detergent lysis protocol described step-by-step in Matson et. al. 2018 [30]. Following sample preparation, the single nuclei suspensions were delivered to collaborators at the NHLBI genomics core facility and processed for single-cell sequencing using the 10X Genomics Chromium Single Cell 3’ Kit. Samples were sequenced using the Illumina, Inc. NovaSeq 6000 system to yield a single nuclei data set. 10x Genomics Cellranger 3.0.0 was used to map sequences to a reference mouse genome [136].

#### Clustering Analysis

The data were analyzed using the R package Seurat version 4.0 developed by the Satija Lab [36]. Clustering was performed in three phases on (1) all cell types, (2) all neurons, and (3) putative PVT neurons, based on the methods described in Russ et. al. 2021 with the following adaptations [137]. For all three phases, each cluster was analyzed for candidate marker genes and excluded if the cluster met either of the following criteria. Clusters were considered low-quality if they had fewer than three significant markers relevant to cell type, particularly if they showed very low nGene (<100). Clusters were considered doublets if they had significant markers for multiple unrelated cell types and boxplots indicate they had a significant range of nGene. Differential gene analysis was performed using the Wilcoxon Rank Sum test with log fold change >0.5 and p value adjusted<0.05.

For phase one, data from all samples were pooled together and filtered based on the following criteria: nuclei containing less than 200 genes (to filter empty droplets), nuclei with greater than 5% mitochondrial genes were removed (to filter low quality nuclei). The data set was normalized using the SCTransform package and highest variable genes were used to perform principal component analysis (PCA). The Harmony algorithm was used to penalize clusters biased by sample origin [31]. The most significant principal components were identified by elbow plot and manual inspection of the contributing gene lists and 13 PCs were used for clustering. To select cluster resolution, a range of values were tested from 0.1-0.8 and UMAPs and top differentially expressed gene lists were generated and visually inspected, and resolution 0.5 was selected. Nuclei were clustered and visualized by using UMAP. Cell types were classified using DropViz and based on the presence of well-established marker genes (http://dropviz.org) [138]. For phase 2, raw data from all cells in neuronal clusters were used, filtered for nuclei containing at least 1,000 detected genes, re-scaled, re-normalized, and re-integrated. The top 6 PCs were selected and resolution 0.3 was selected, using the approach described above. Here, neurons were then classified based on (1) comparison of cluster markers with known markers and with known co-expression patterns in the literature or Allen Brain In Situ Hybridization Atlas (http://mouse.brain-map.org). For phase 3, targeted sub-clustering was performed to investigate PVT specific subtypes and an additional subtype was identified that did not resolve in previous phases. The above procedure was repeated and 6PCs and resolution 0.25 were used. Top marker genes were selected for each PVT cluster by calculating the ratio of expression for a particular gene across the active cluster (pct. 1) and all other clusters (pct. 2) and selecting for the highest ratio value (pct. ratio) (Supp. Table 3). Higher pct. ratio values indicate markers with higher specificity to a given cluster. *Esr1* (PVT1; pct. ratio = 2.031), *Col12a1* (PVT2; pct. ratio = 4.648), *Gal* (PVT3; pct. ratio = 2.358), *Hcrtr1* (PVT4; pct. ratio = 6.190), and *Drd3* (PVT5; pct. ratio = 2.155).

### *In situ* hybridization

#### Sample preparation and ISH procedure (RNAscope)

Fresh-frozen brains from adult male C57BL/6J mice (8-12 weeks) were sectioned at a thickness of 16 µm using a Cryostat (Leica Biosystems). Sections were collected onto Superfrost Plus slides (Daigger Scientific, Inc), immediately placed on dry ice and subsequently transferred to a −80°C freezer. *Esr1*, *Col12a1*, *Gal, Hcrtr1, Drd2*, *Npffr1*, *Npsr1, Chrm2* mRNA signal was labeled by using the Hiplex RNAscope Fluorescent kit v2 (Advanced Cell Diagnostics), according to manufacturer’s instructions. Sections were cover slipped using Diamond Prolong antifade mounting medium with DAPI (ThermoFisher Scientific).

#### Signal detection and analysis

After the amplification procedure, slides were examined on a Nikon A1R HD confocal microscope (Nikon) using a 20X objective. Images were first processed by removing background noise using the background subtraction tool (5.0 pixel rolling ball radius) in FIJI (Image J). Images from the same section were then registered using RNAscope Hiplex Image Registration Software v2. Signal was subsequently quantified with CellProfiler using a freely available pipeline (macros) for RNAscope [139]. A protocol with a step-by-step description of how to implement this pipeline for analyzing RNAscope data was recently published [140]. This pipeline was modified to include up to 12 mRNA probes and cells were considered positive for a given gene if they contained a minimum of 7 mRNA transcripts (dots). Subsequently, all RNAscope data was analyzed by a blind experimenter. Sections from bregma −0.22 to −0.34 were considered anterior PVT and sections from bregma −1.58 to −1.70 were considered posterior PVT. Gene expression matrices were the generated and analyzed using RStudio Version 4.2.0. Heatmaps and hierarchical clustering were performed using R functions pheatmap() and hclust().

### Merged analysis and integration

#### Published data acquisition

Published data from the ThalamoSeq project were downloaded from the NCBI Gene Expression Omnibus public functional genomics data repository. Single-cell data and metadata from Phillips et. al. 2019 (GSE133911) was downloaded as processed count matrices (https://www.ncbi.nlm.nih.gov/geo/query/acc.cgi?acc=GSE133911) [29].

#### RPCA-based integration and Clustering Analysis

Count matrices for each dataset were merged to obtain the full data file. Uniform data filtering was applied across the merged file. All cells and nuclei with at least 1,000 detected genes (to exclude low quality neurons) and less than 5% of transcripts being mitochondrial (to exclude lysing cells or mitochondria-nuclei doublets) were analyzed, yielding a total of 8,507 nuclei.

The merged data were analyzed using Seurat version 4.0 [36]. The integration was performed using Seurat version 4.0 Standard Workflow (RPCA) Integration such that data are first normalized with SCTransform (method = “glmGamPoi”) and PCA is run individually on each dataset prior to integration. Integration anchors were calculated using 30 PCs and used to integrate the data.

The most significant principal components were identified by elbow plot and manual inspection of the contributing gene lists and 13 PCs were used for clustering. To select cluster resolution, a range of values were tested from 0.2-0.6 and UMAPs and top differentially expressed gene lists were generated, and resolution 0.35 was selected. Conserved markers for integrated data cluster 1 were obtained using FindConservedMarkers() function to confirm that these PVT/Prefrontal-projectors express both *Col12a1* and *Chrm2*.

### Statistics and data presentation

All data were imported to OriginPro 2016 (OriginLab Corp.) for statistical analyses. Initially, normality tests (D’Agostino-Pearson and Kolmogorov-Smirnov) were performed to determine the appropriate of the statistical tests used. All data are presented as mean ± s.e.m. No assumptions or corrections were made prior to data analysis. Differences between two groups were always examined using a two-sided Student’s t-test, where *P*<0.05 was considered significant and *P*>0.05 was considered non-significant. The sample sizes used in our study are about the same or exceed those estimated by power analysis (power = 0.9, α = 0.05). Graphs were generated in OriginPro or Rstudio and figures were generated using Adobe Illustrator. For RNAscope experiments the sample size is 2-3 mice. All experiments were replicated at least once, and all subjects were age matched.

## Data availability

All the data that support the findings presented in this study are available from the corresponding author upon reasonable request. Raw and processed RNA-seq data have been deposited into the Gene Expression Omnibus repository (pending).

## ACKNOWLEDGEMENTS

We thank the NHLBI Genomics Core for performing library preparation and sequencing our samples. We thank Drs. Sofia Beas and Hugo Tejeda for their comments on the manuscript. We thank Drs. Ruchi Komal, Maria Yurgel, Anton Schulmann, and Jesse Zhan for their training and guidance. This work was supported by the NIMH Intramural Research Program (1ZIAMH002950) and (in part) by the Division of Intramural Research of the NIH, NINDS (ZIANS003153). The content is solely the responsibility of the author(s) and does not necessarily represent the official views of the National Institutes of Health.

## SUPPLEMENTARY FIGURES WITH LEGENDS

**Figure 1—figure supplement 1.**
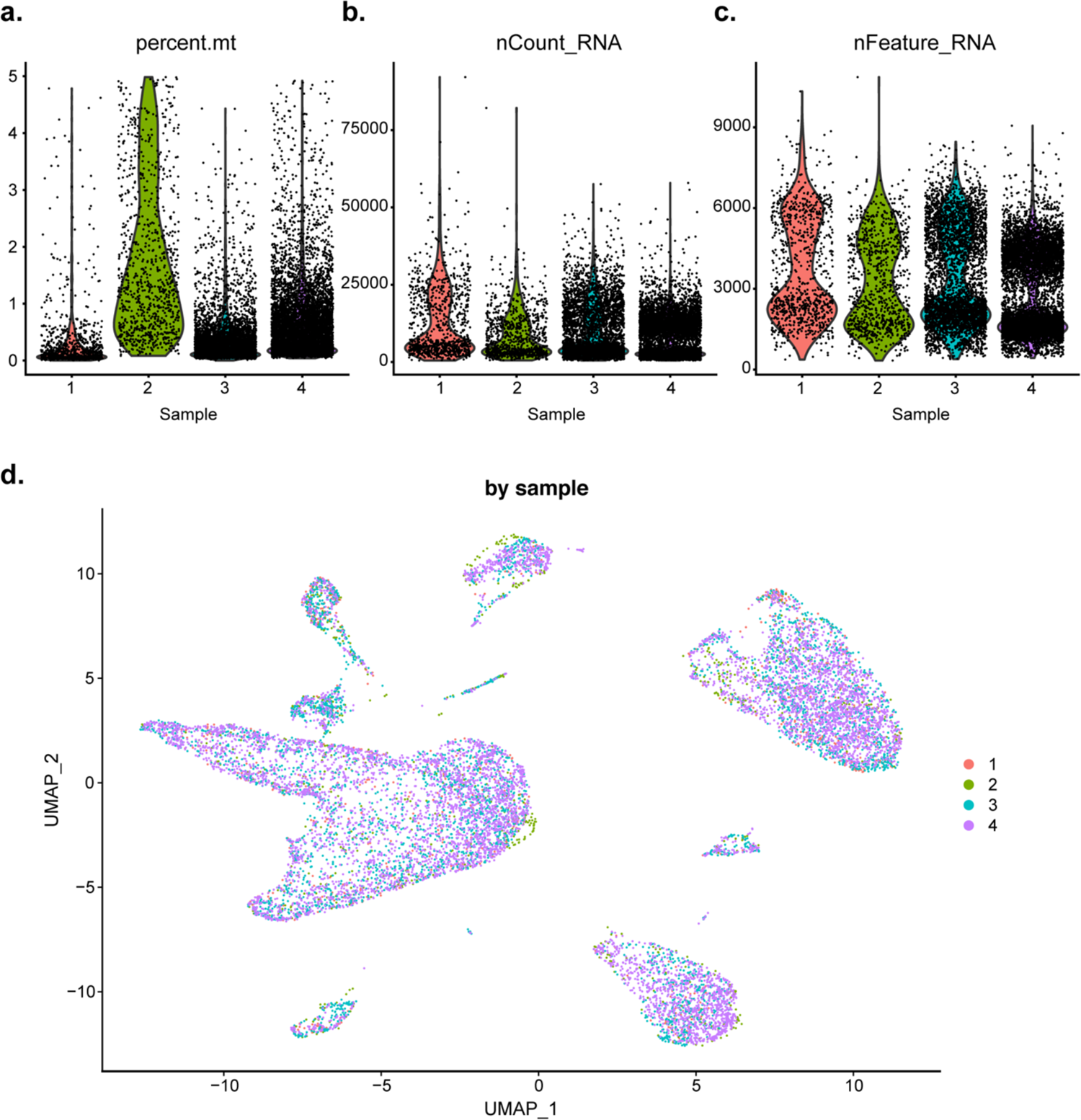
Quality control statistics and UMAP by sample. **a-c.** Violin plots depicting the percent mitochondrial RNA (a), number of UMIs (b), and number of genes (c) detected in each of the four samples. **d.** The UMAP plot of the same nuclei from Figure 1c labeled by sample of origin.

**Figure 1—figure supplement 2.**
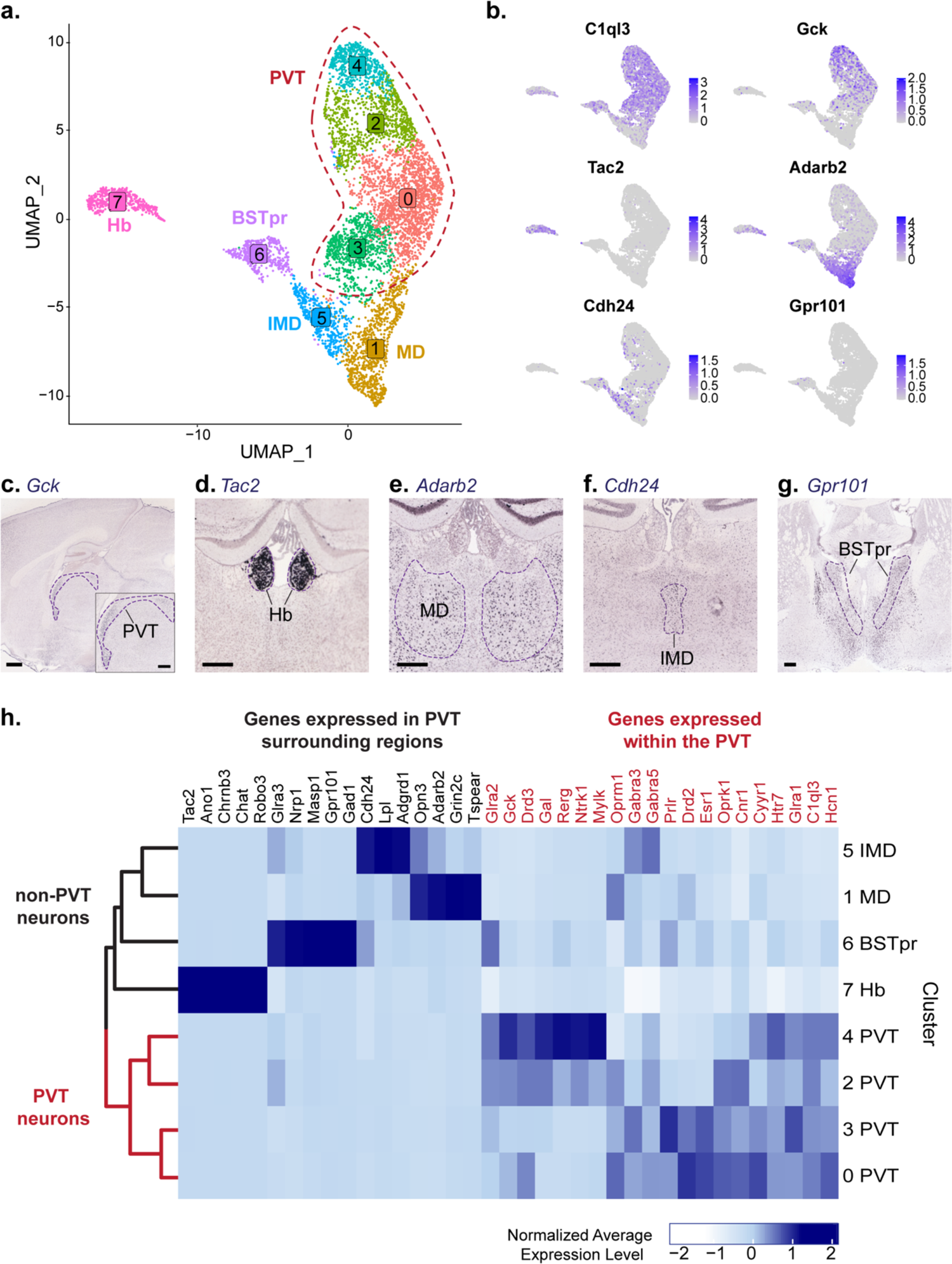
Clustering analysis and classification of all neurons. **a.** The UMAP plot of 6,555 neuronal nuclei shows 8 clusters. **b.** Feature plots of representative gene expression of markers from different clusters. **c.** In situ hybridization of *Gck* (ABA Experiment #68269269), scale bar 500um, outline depicts the PVT. **d.** In situ hybridization of *Tac2* (ABA Experiment #72339556), scale bar 300um, outline depicts the Hb. **e.** In situ hybridization of *Adarb2* (ABA Experiment #73925721), scale bar 300um, outline depicts the MD. **f.** In situ hybridization of *Cdh24* (ABA Experiment #70231307), scale bar 300um, outline depicts the IMD. **g.** In situ hybridization *of Gpr101* (ABA Experiment #79591639), scale bar 300um, outline depicts the BSTpr. **h.** Heatmap of normalized average gene expression of PVT-enriched genes (red) and PVT-depleted genes (black). ABA, Allen Brain Atlas.

**Figure 2—figure supplement 1.**
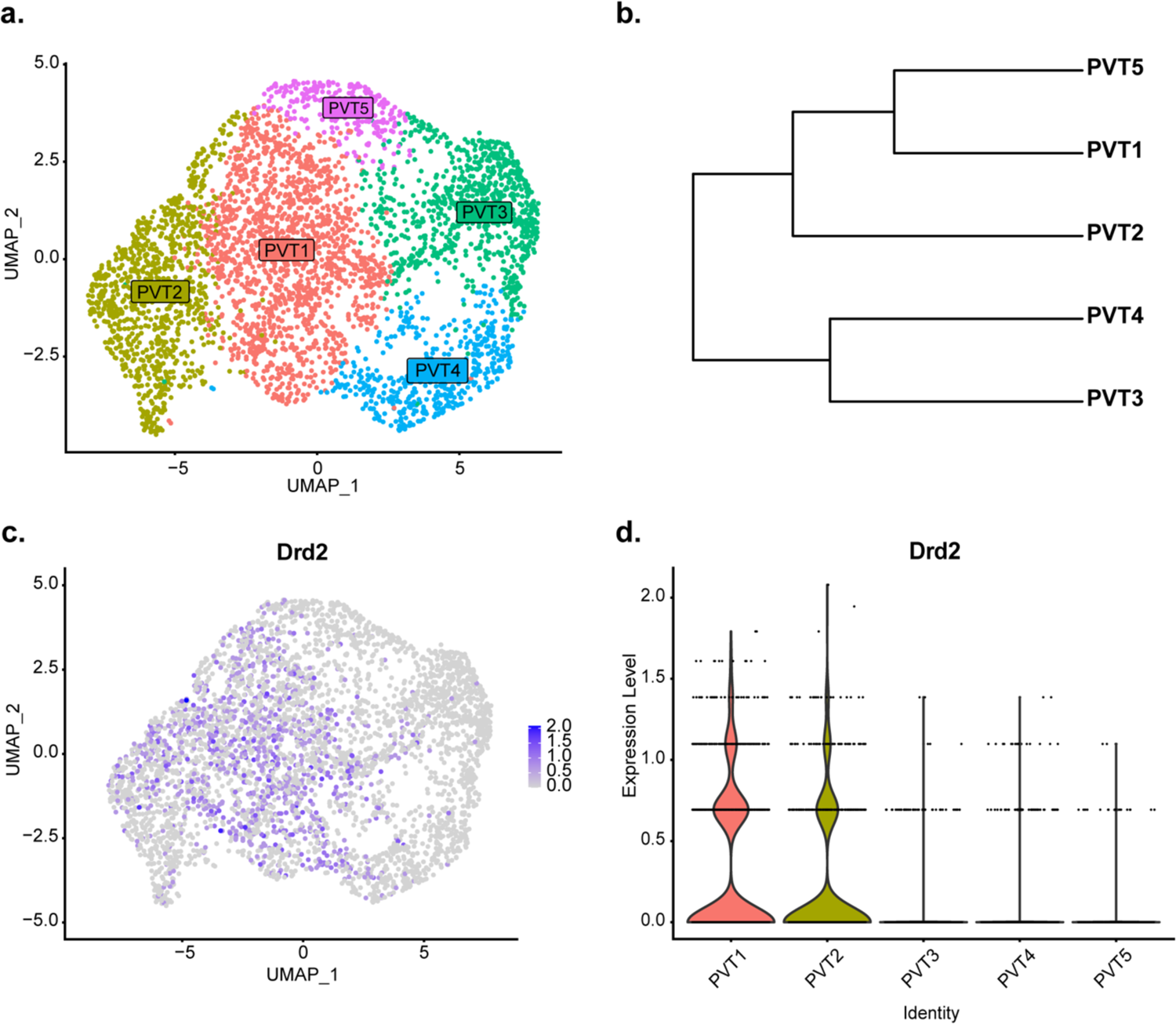
*Drd2* expression is found in PVT1 and PVT2. **a.** The UMAP plot of 4,067 PVT neuronal nuclei shows 5 clusters. **b.** Phylogenetic tree depicting cluster relationships based on distance between clusters in gene expression space. c. Feature plot of *Drd2* expression across all PVT neurons. d. Violin plot of *Drd2* expression level across 5 PVT clusters.

**Figure 3—figure supplement 1.**
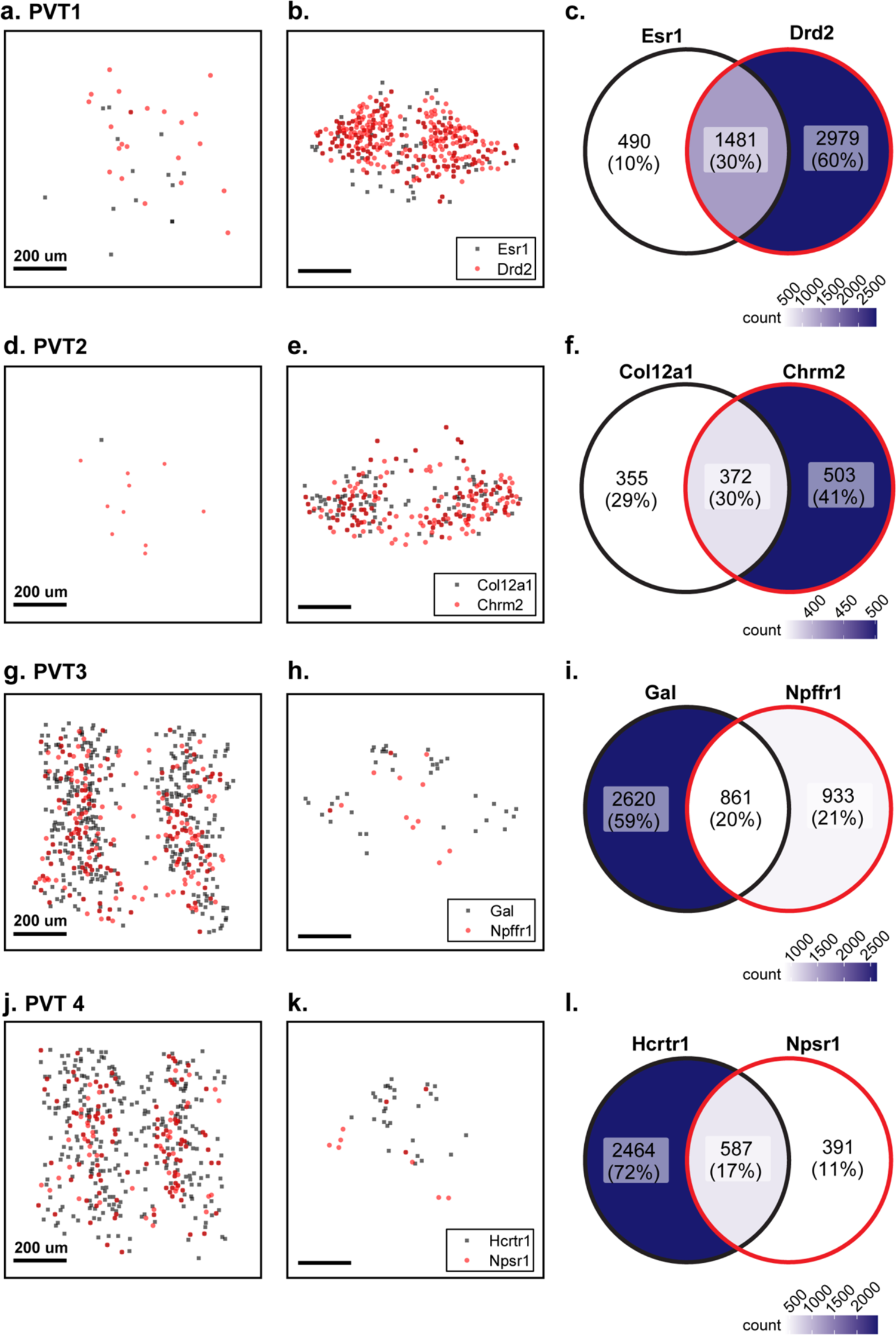
Spatial distribution and overlap across anterior- or posterior- and pan-PVT subtypes. **a.** Dendrogram depicting Euclidian distance between cluster types based on gene transcript expression. **b-c.** Location of positive cells expressing *Esr1* (blue square), *Col12a1* (black circle), and *Drd3* (yellow triangle) in aPVT (b) or pPVT (c). **d.** Venn diagram of number and percent of *Esr1*-, *Col12a1*-, or *Drd3*-positive cells. **e-f.** Location of positive cells expressing *Gal* (green square), *Hcrtr1* (magenta circle), or *Drd3* (yellow triangle) in aPVT (e) or pPVT (f). **g.** Venn diagram of number and percent of *Gal*-, *Hcrtr1*-, or *Drd3*-positive cells. Venn Diagram legends, color intensity correlates to cell count.

**Figure 3—figure supplement 2.**
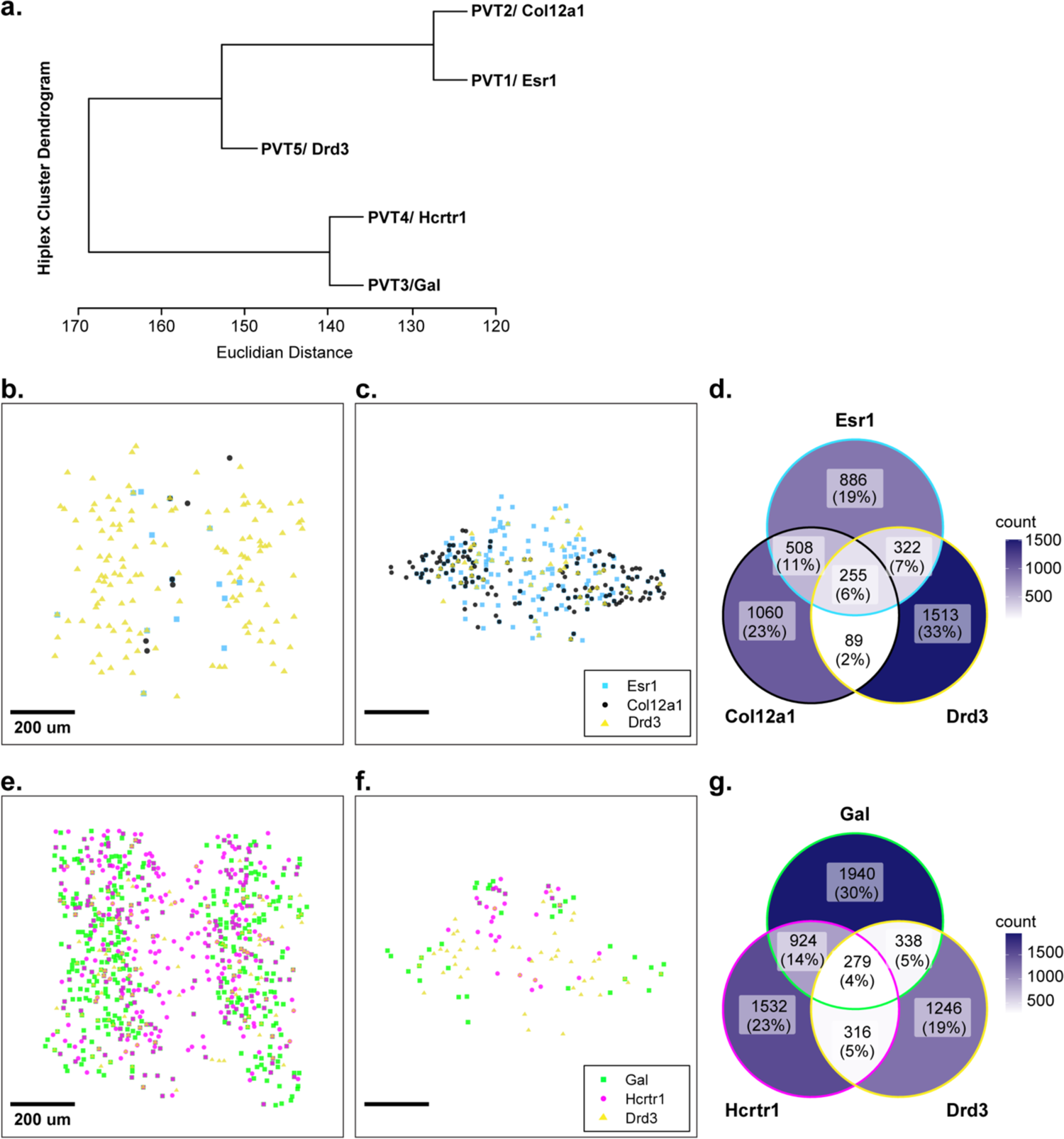
Overlap in gene expression among top markers from the same cluster. **a-b.** Location of *Esr1* (black) or *Drd2* (red) positive cells in aPVT (a) or pPVT (b) from PVT1 subtype. **c.** Venn diagram of cell count and percent overlap between *Esr1*- and *Drd2*-positive cells. **d-e.** Location of *Col12a1* (black) or *Chrm2* (red) positive cells in aPVT (a) or pPVT (b) from PVT2 subtype. **f.** Venn diagram of cell count and percent overlap between *Col12a1*- and *Chrm2*-positive cells. **g-i.** Location of *Gal* (black) or *Npffr1* (red) positive cells in aPVT (a) or pPVT (b) from PVT3 subtype. **f.** Venn diagram of cell count and percent overlap between *Gal*- and *Npffr1*-positive cells. **j-k.** Location of *Hcrtr1* (black) or *Npsr1* (red) positive cells in aPVT (a) or pPVT (b) from PVT4 subtype. **f.** Venn diagram of cell count and percent overlap between *Hcrtr1*- and *Npsr1*-positive cells. Venn Diagram legends, color intensity correlates to cell count. Black scale bar indicates 200um.

**Figure 3—figure supplement 3.**
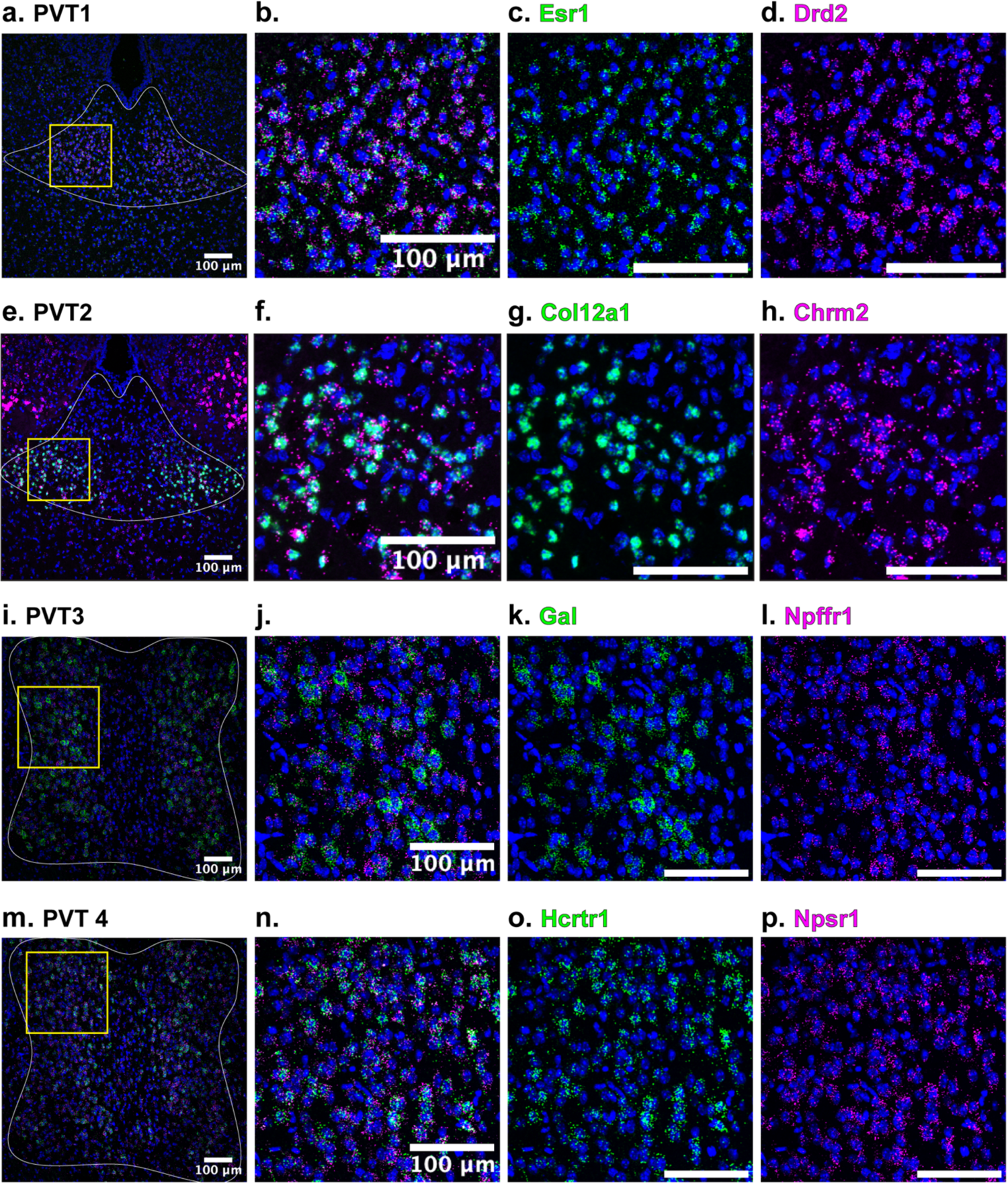
In situ hybridization of top markers from the same cluster. **a.** In situ hybridization of *Esr1* (green), *Drd2* (magenta), and DAPI (blue) in pPVT (white outline). **b-d.** Yellow square insert from (a). **e.** In situ hybridization of *Col12a1* (green), *Chrm2* (magenta), and DAPI (blue) in pPVT (white outline). **f-d.** Yellow square insert from (e). **i.** In situ hybridization of *Gal* (green), *Npffr1* (magenta), and DAPI (blue) in aPVT (white outline). **j-l.** Yellow square insert from (i). **m.** In situ hybridization of *Hcrtr1* (green), *Npsr1* (magenta), and DAPI (blue) in aPVT (white outline). **n-p.** Yellow square insert from (m). All images are 20x representative confocal images with brightness and contrast adjusted depicting expression patterns found in all sections from N=3 animals. White scale bar indicates 100um.

**Figure 5—figure supplement 1.**
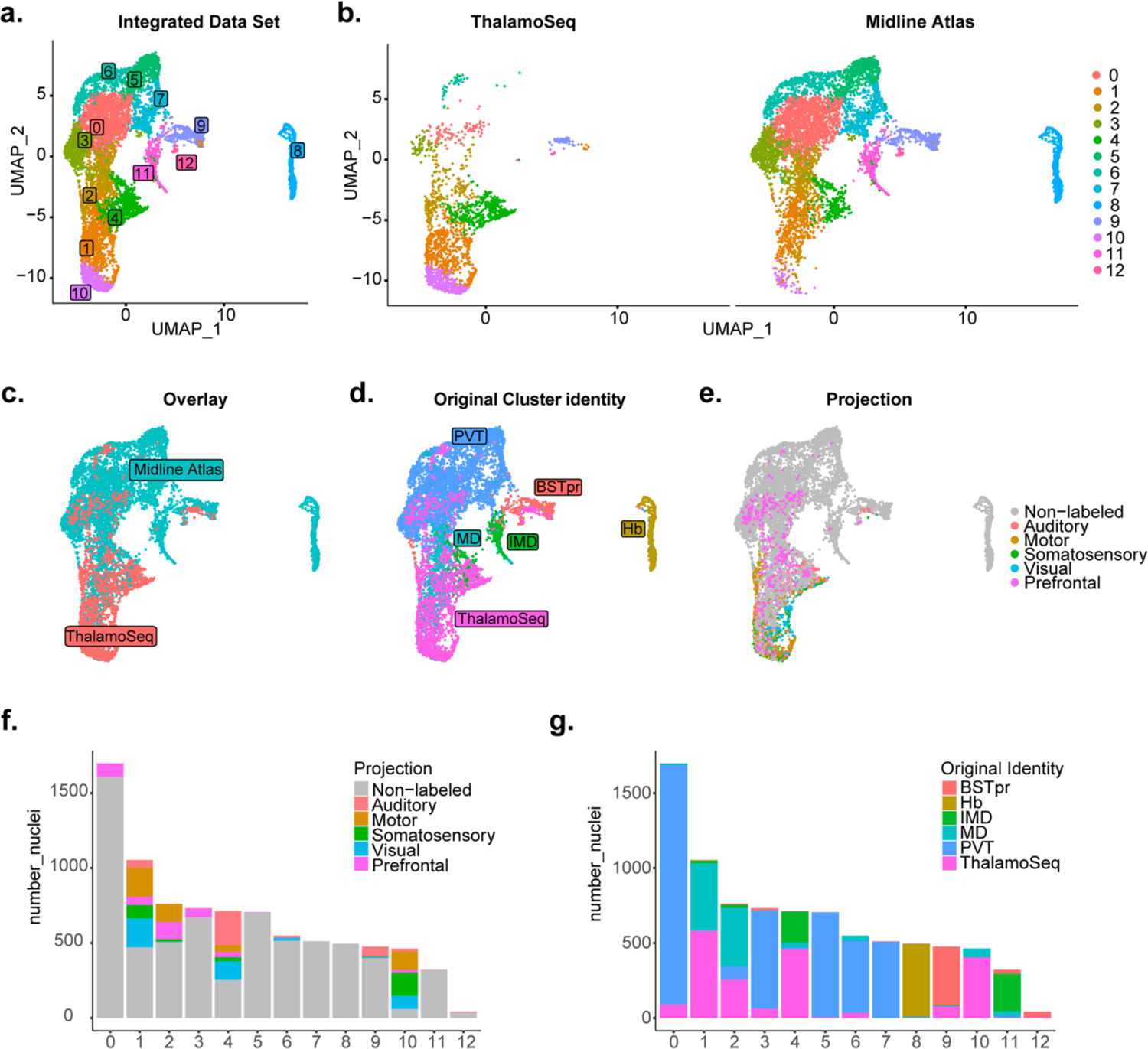
RPCA-based integration with Thalamoseq dataset and all neurons from present study reveals mostly non-overlapping transcriptomes. **a.** The UMAP plot of 8,527 cells or nuclei from combined datasets annotated by cluster identity. **b.** The same cells/nuclei separated by study of origin: Thalamoseq (left) and present study (right) and overlayed in **c. d.** The same cells/nuclei colored by original identity. **e.** The same cells/nuclei colored by cortical projection target. **f.** Proportion of nuclei in each cluster labeled by cortical projection target. **g.** Proportion of nuclei in each cluster labeled by original identity.

**Figure 5—figure supplement 2.**
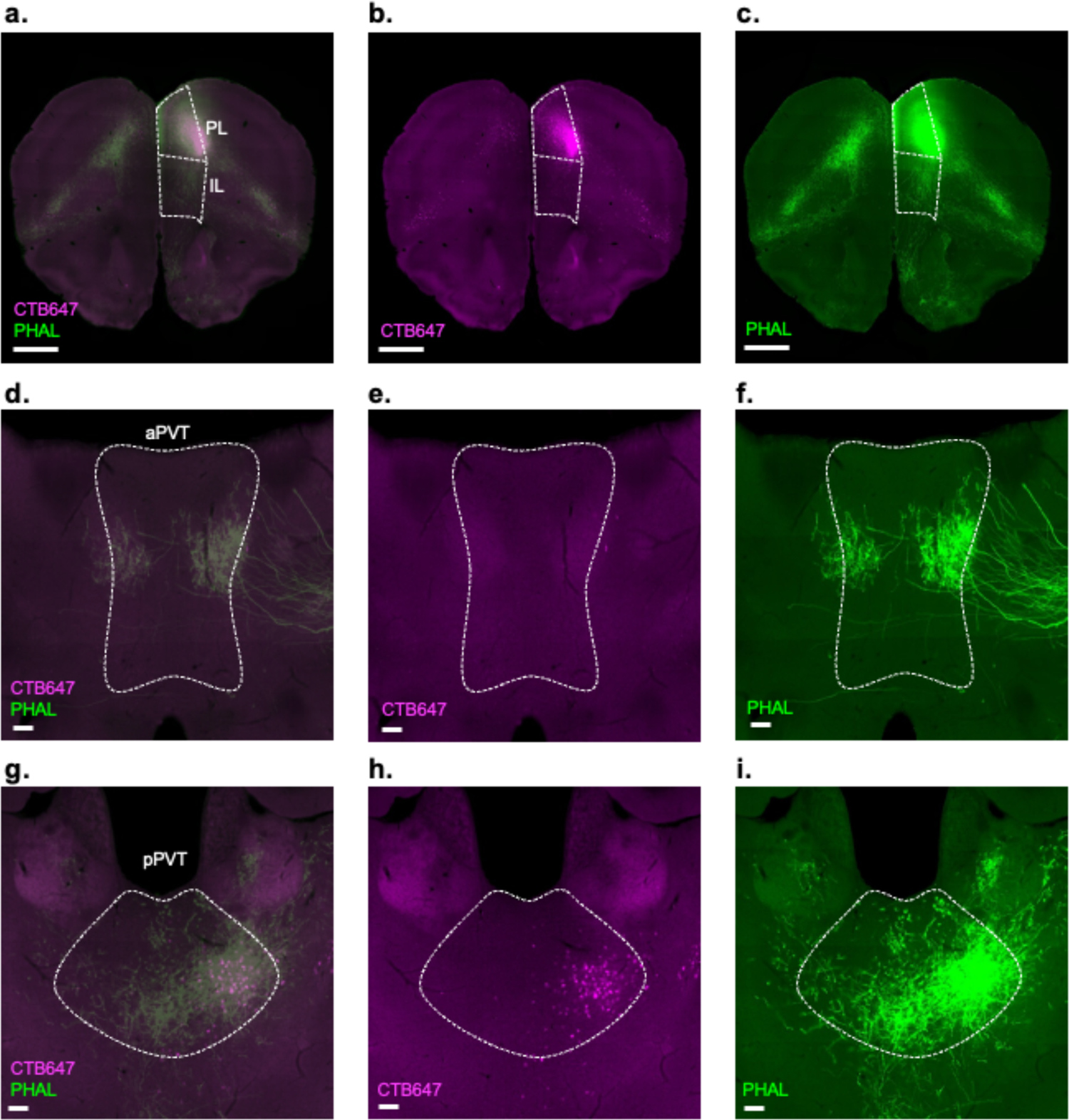
Dual anterograde and retrograde tracing from the prelimbic cortex (Mouse Connectome Project Experiment SW120125-02A) **a-c.** Representative image of the injection site in the prelimbic cortex (PL) of retrograde tracer CTB647 (magenta) and anterograde tracer PHAL (green); Scale bar 1mm. **d-f.** Representative image of retrogradely labeled cells (magenta) and anterogradely labeled fibers (green) in the aPVT; Scale bar 100um. **g-i.** Representative image of retrogradely labeled cells (magenta) and anterogradely labeled fibers (green) in the pPVT; Scale bar 100um. Legend: IL, infralimbic cortex; www.mouseconnectome.org

**Figure 5—figure supplement 3.**
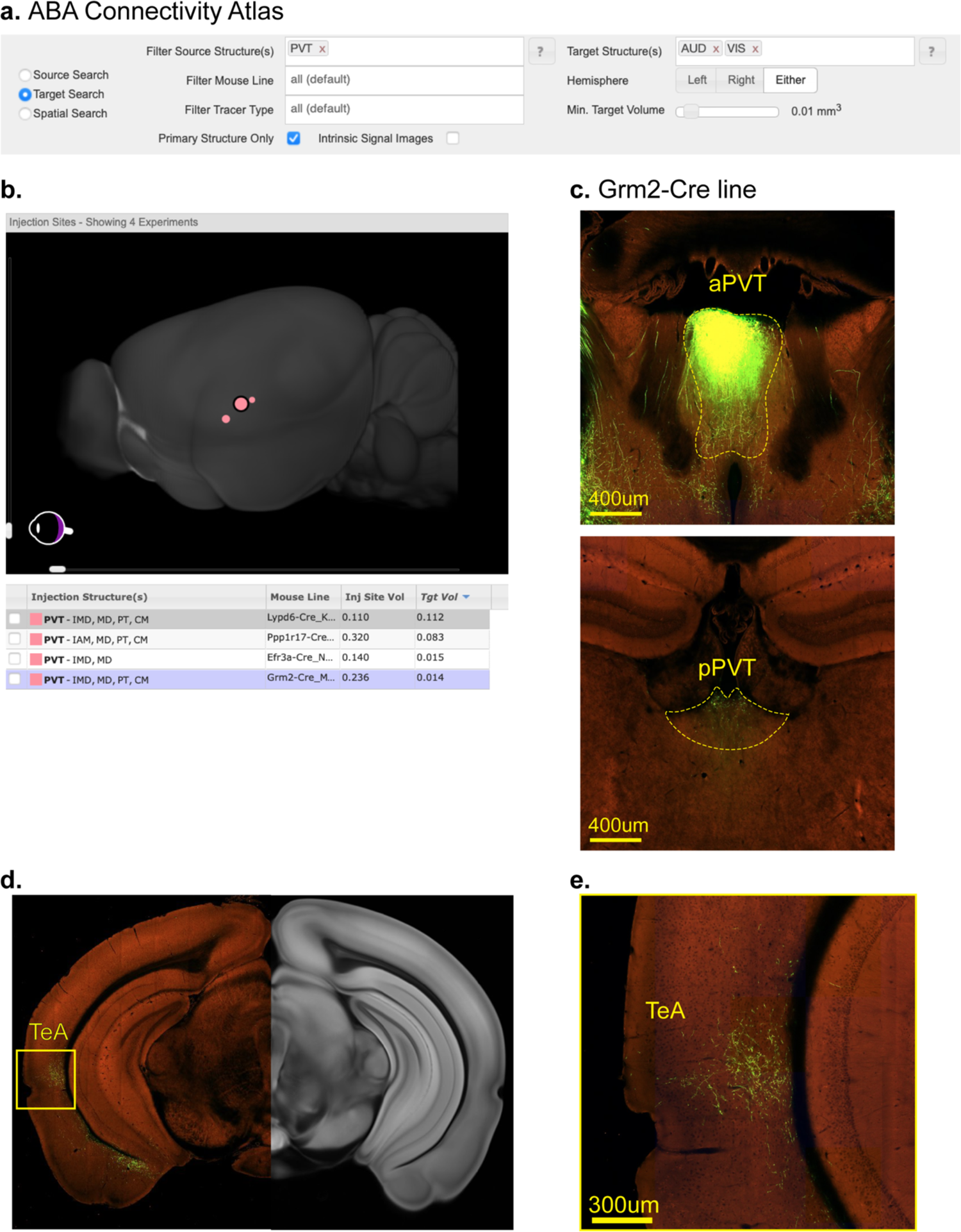
Projections from aPVT to Auditory and Visual cortical areas. **a.** Schematic of Target Search tool used to identify experiments where projections from the PVT to Auditory (AUD) and Visual (VIS) areas were detected (Mouse Brain Connectivity Atlas of the Allen Brian Institute, https://connectivity.brain-map.org). **b.** Four experiments identified from target search. **c.** Representative image from Experiment #183225830 depicting the injection site in the aPVT (top) and pPVT (bottom). **d.** Representative image of fibers located in the temporal association cortex (TeA) from Experiment #183225830 **e.** Same image of yellow insert from (d).

**Figure 5—figure supplement 4.**
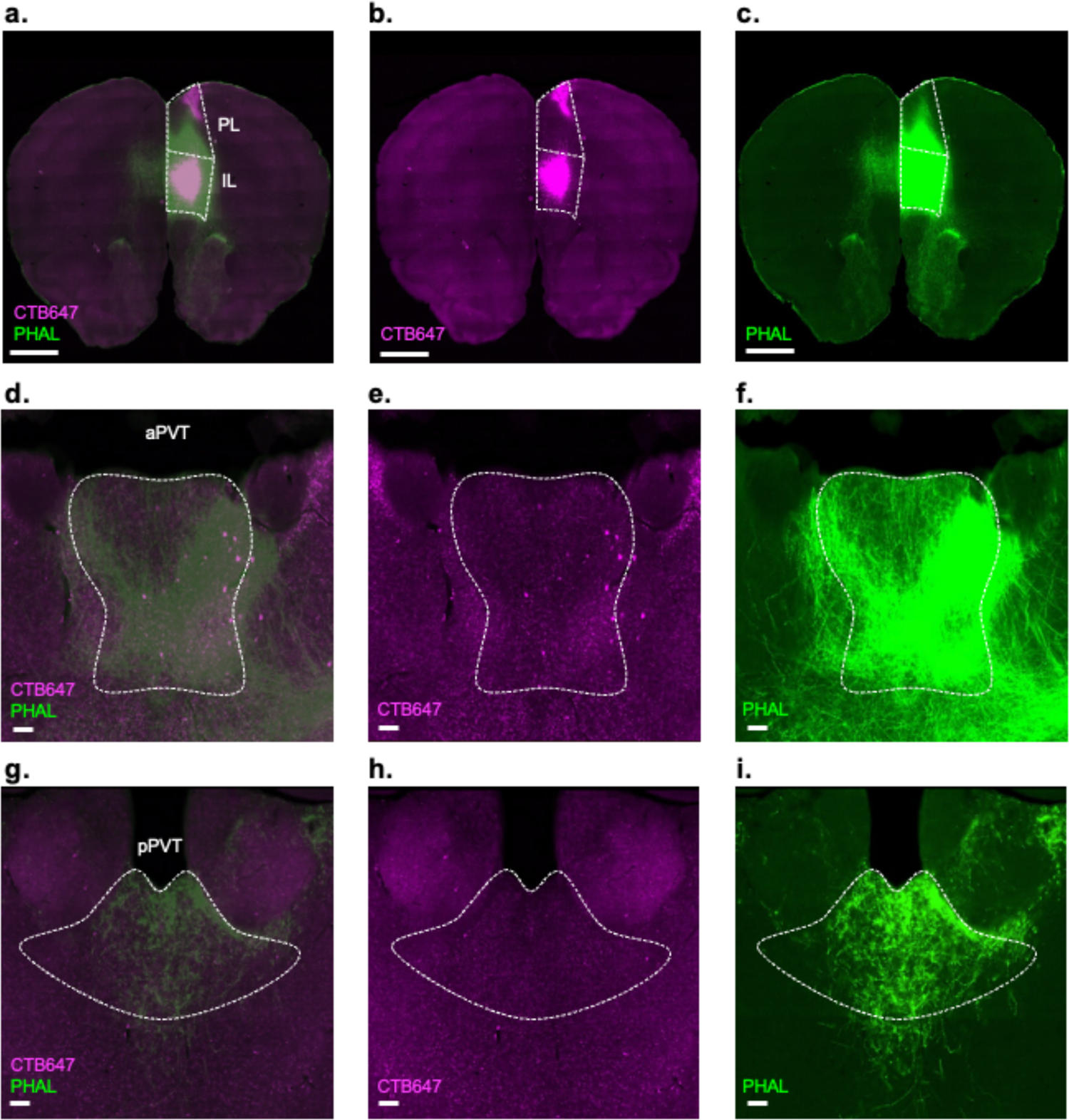
Dual anterograde and retrograde tracing from the infralimbic cortex (Mouse Connectome Project Experiment SW110614-04B) **a-c.** Representative image of the injection site in the infralimbic cortex (IL) of retrograde tracer CTB647 (magenta) and anterograde tracer PHAL (green); Scale bar 1mm. **d-f.** Representative image of retrogradely labeled cells (magenta) and anterogradely labeled fibers (green) in the aPVT; Scale bar 100um. **g-i.** Representative image of retrogradely labeled cells (magenta) and anterogradely labeled fibers (green) in the pPVT; Scale bar 100um; Legend: PL, prelimbic cortex; www.mouseconnectome.org

## AUTHOR CONTRIBUTIONS

C.G and Y.L performed all experiments in this study. C.G, and M.A.P. designed the study. A.J.L. provided critical feedback on the acquisition and analysis of snRNA-Seq data. C.G and C. A. G. analyzed the data. C.G. and M.A.P. interpreted results and wrote the paper.

## REFERENCES

1. Petrovich, G.D., The Function of Paraventricular Thalamic Circuitry in Adaptive Control of Feeding Behavior. Front Behav Neurosci, 2021. 15: p. 671096.

2. Kirouac, G.J., The Paraventricular Nucleus of the Thalamus as an Integrating and Relay Node in the Brain Anxiety Network. Front Behav Neurosci, 2021. 15: p. 627633.

3. McNally, G.P., Motivational competition and the paraventricular thalamus. Neurosci Biobehav Rev, 2021. 125: p. 193–207.

4. Penzo, M.A. and C. Gao, The paraventricular nucleus of the thalamus: an integrative node underlying homeostatic behavior. Trends Neurosci, 2021. 44(7): p. 538–549.

5. Kelley, A.E., B.A. Baldo, and W.E. Pratt, A proposed hypothalamic-thalamic-striatal axis for the integration of energy balance, arousal, and food reward. J Comp Neurol, 2005. 493(1): p. 72–85.

6. Barson, J.R., N.R. Mack, and W.-J. Gao, The Paraventricular Nucleus of the Thalamus Is an Important Node in the Emotional Processing Network. Frontiers in Behavioral Neuroscience, 2020. 14(191).

7. McGinty, J.F. and J.M. Otis, Heterogeneity in the Paraventricular Thalamus: The Traffic Light of Motivated Behaviors. Frontiers in Behavioral Neuroscience, 2020. 14(181).

8. Matyas, F., et al., A highly collateralized thalamic cell type with arousal-predicting activity serves as a key hub for graded state transitions in the forebrain. Nature Neuroscience, 2018. 21(11): p. 1551-+.

9. Hua, R., et al., Calretinin Neurons in the Midline Thalamus Modulate Starvation-Induced Arousal. Curr Biol, 2018. 28(24): p. 3948–3959 e4.

10. Gao, C., et al., Two genetically, anatomically and functionally distinct cell types segregate across anteroposterior axis of paraventricular thalamus. Nat Neurosci, 2020. 23(2): p. 217–228.

11. Bhatnagar, S. and M.F. Dallman, The paraventricular nucleus of the thalamus alters rhythms in core temperature and energy balance in a state-dependent manner. Brain Res, 1999. 851(1-2): p. 66–75.

12. Stratford, T.R. and D. Wirtshafter, Injections of muscimol into the paraventricular thalamic nucleus, but not mediodorsal thalamic nuclei, induce feeding in rats. Brain Res, 2013. 1490: p. 128–33.

13. Labouebe, G., et al., Glucose-responsive neurons of the paraventricular thalamus control sucrose-seeking behavior. Nat Neurosci, 2016. 19(8): p. 999–1002.

14. Sofia Beas, B., et al., A ventrolateral medulla-midline thalamic circuit for hypoglycemic feeding. Nature Communications, 2020. 11(1): p. 6218.

15. Meffre, J., et al., Orexin in the Posterior Paraventricular Thalamus Mediates Hunger-Related Signals in the Nucleus Accumbens Core. Curr Biol, 2019. 29(19): p. 3298–3306 e4.

16. Iglesias, A.G. and S.B. Flagel, The Paraventricular Thalamus as a Critical Node of Motivated Behavior via the Hypothalamic-Thalamic-Striatal Circuit. Frontiers in Integrative Neuroscience, 2021. 15.

17. Li, S. and G.J. Kirouac, Projections from the paraventricular nucleus of the thalamus to the forebrain, with special emphasis on the extended amygdala. J Comp Neurol, 2008. 506(2): p. 263–87.

18. Vertes, R.P. and W.B. Hoover, Projections of the paraventricular and paratenial nuclei of the dorsal midline thalamus in the rat. J Comp Neurol, 2008. 508(2): p. 212–37.

19. Moga, M.M., R.P. Weis, and R.Y. Moore, Efferent projections of the paraventricular thalamic nucleus in the rat. J Comp Neurol, 1995. 359(2): p. 221–38.

20. Kirouac, G.J., Placing the paraventricular nucleus of the thalamus within the brain circuits that control behavior. Neurosci Biobehav Rev, 2015. 56: p. 315–29.

21. Huang, Z.J. and H. Zeng, Genetic approaches to neural circuits in the mouse. Annu Rev Neurosci, 2013. 36: p. 183–215.

22. Huang, Z.J., Toward a genetic dissection of cortical circuits in the mouse. Neuron, 2014. 83(6): p. 1284–1302.

23. Fuzik, J., et al., Integration of electrophysiological recordings with single-cell RNA-seq data identifies neuronal subtypes. Nat Biotechnol, 2016. 34(2): p. 175–183.

24. Oh, S.W., et al., A mesoscale connectome of the mouse brain. Nature, 2014. 508(7495): p. 207-214.

25. Luo, L., E.M. Callaway, and K. Svoboda, Genetic dissection of neural circuits. Neuron, 2008. 57(5): p. 634–60.

26. Luo, L., E.M. Callaway, and K. Svoboda, Genetic Dissection of Neural Circuits: A Decade of Progress. Neuron, 2018. 98(2): p. 256–281.

27. Kessler, S., et al., Glucokinase neurons of the paraventricular nucleus of the thalamus sense glucose and decrease food consumption. iScience, 2021. 24(10): p. 103122.

28. Engelke, D.S., et al., A hypothalamic-thalamostriatal circuit that controls approach-avoidance conflict in rats. Nature Communications, 2021. 12(1): p. 2517.

29. Phillips, J.W., et al., A repeated molecular architecture across thalamic pathways. Nat Neurosci, 2019.

30. Au - Matson, K.J.E., et al., Isolation of Adult Spinal Cord Nuclei for Massively Parallel Single-nucleus RNA Sequencing. JoVE, 2018(140): p. e58413.

31. Korsunsky, I., et al., Fast, sensitive and accurate integration of single-cell data with Harmony. Nature Methods, 2019. 16(12): p. 1289–1296.

32. Lein, E.S., et al., Genome-wide atlas of gene expression in the adult mouse brain. Nature, 2007. 445(7124): p. 168-176.

33. Gaspari, S., et al., Structural and molecular characterization of paraventricular thalamic glucokinase-expressing neuronal circuits in the mouse. J Comp Neurol, 2022. 530(11): p. 1773–1949.

34. Chew, K.S., et al., Anatomical and Behavioral Investigation of C1ql3 in the Mouse Suprachiasmatic Nucleus. J Biol Rhythms, 2017. 32(3): p. 222–236.

35. Kolaj, M., et al., Midline thalamic paraventricular nucleus neurons display diurnal variation in resting membrane potentials, conductances, and firing patterns in vitro. J Neurophysiol, 2012. 107(7): p. 1835–44.

36. Hao, Y., et al., Integrated analysis of multimodal single-cell data. Cell, 2021. 184(13): p. 3573–3587.e29.

37. Ståhl, P.L., et al., Visualization and analysis of gene expression in tissue sections by spatial transcriptomics. Science, 2016. 353(6294): p. 78-82.

38. Tasic, B., et al., Shared and distinct transcriptomic cell types across neocortical areas. Nature, 2018. 563(7729): p. 72-78.

39. Cembrowski, M.S. and N. Spruston, Heterogeneity within classical cell types is the rule: lessons from hippocampal pyramidal neurons. Nat Rev Neurosci, 2019. 20(4): p. 193–204.

40. Romanov, R.A., et al., Unified Classification of Molecular, Network, and Endocrine Features of Hypothalamic Neurons. Annual Review of Neuroscience, 2019. 42(1): p. 1–26.

41. O’Leary, T.P., et al., Extensive and spatially variable within-cell-type heterogeneity across the basolateral amygdala. eLife, 2020. 9: p. e59003.

42. Moffitt, J.R., et al., Molecular, spatial, and functional single-cell profiling of the hypothalamic preoptic region. Science, 2018. 362(6416).

43. Li, Y., et al., Distinct subnetworks of the thalamic reticular nucleus. Nature, 2020. 583(7818): p. 819-824.

44. Mukherjee, A., et al., Thalamic circuits for independent control of prefrontal signal and noise. Nature, 2021. 600(7887): p. 100-104.

45. Martinez-Garcia, R.I., et al., Two dynamically distinct circuits drive inhibition in the sensory thalamus. Nature, 2020. 583(7818): p. 813-818.

46. Roy, D.S., et al., Thalamic subnetworks as units of function. Nature Neuroscience, 2022. 25(2): p. 140–153.

47. Krout, K.E. and A.D. Loewy, Parabrachial nucleus projections to midline and intralaminar thalamic nuclei of the rat. Journal of Comparative Neurology, 2000. 428(3): p. 475–494.

48. Kirouac, G.J., M.P. Parsons, and S. Li, Orexin (hypocretin) innervation of the paraventricular nucleus of the thalamus. Brain Res, 2005. 1059(2): p. 179–88.

49. Kirouac, G.J., M.P. Parsons, and S. Li, Innervation of the paraventricular nucleus of the thalamus from cocaine- and amphetamine-regulated transcript (CART) containing neurons of the hypothalamus. J Comp Neurol, 2006. 497(2): p. 155–65.

50. García-Cabezas, M.Á., et al., Distribution of the dopamine innervation in the macaque and human thalamus. NeuroImage, 2007. 34(3): p. 965–984.

51. Freedman, L.J. and M.D. Cassell, Relationship of thalamic basal forebrain projection neurons to the peptidergic innervation of the midline thalamus. J Comp Neurol, 1994. 348(3): p. 321–42.

52. Vertes, R.P., S.B. Linley, and W.B. Hoover, Pattern of distribution of serotonergic fibers to the thalamus of the rat. Brain Struct Funct, 2010. 215(1): p. 1–28.

53. Curtis, G.R., K. Oakes, and J.R. Barson, Expression and Distribution of Neuropeptide-Expressing Cells Throughout the Rodent Paraventricular Nucleus of the Thalamus. Frontiers in Behavioral Neuroscience, 2021. 14.

54. Pandey, S., et al., Neurotensin in the posterior thalamic paraventricular nucleus: inhibitor of pharmacologically relevant ethanol drinking. Addiction biology, 2019. 24(1): p. 3–16.

55. Barson, J.R., H.T. Ho, and S.F. Leibowitz, Anterior thalamic paraventricular nucleus is involved in intermittent access ethanol drinking: role of orexin receptor 2. Addict Biol, 2015. 20(3): p. 469–81.

56. Chen, Z., et al., Dynorphin activation of kappa opioid receptor reduces neuronal excitability in the paraventricular nucleus of mouse thalamus. Neuropharmacology, 2015. 97: p. 259–69.

57. Barson, J.R., et al., Substance P in the anterior thalamic paraventricular nucleus: promotion of ethanol drinking in response to orexin from the hypothalamus. Addict Biol, 2017. 22(1): p. 58–69.

58. Barrett, L.R., J. Nunez, and X. Zhang, Oxytocin activation of paraventricular thalamic neurons promotes feeding motivation to attenuate stress-induced hypophagia. Neuropsychopharmacology, 2021. 46(5): p. 1045–1056.

59. Matzeu, A., et al., Orexin-A/Hypocretin-1 Mediates Cocaine-Seeking Behavior in the Posterior Paraventricular Nucleus of the Thalamus via Orexin/Hypocretin Receptor-2. J Pharmacol Exp Ther, 2016. 359(2): p. 273–279.

60. Clark, A.M., et al., Dopamine D2 Receptors in the Paraventricular Thalamus Attenuate Cocaine Locomotor Sensitization. eNeuro, 2017. 4(5).

61. Beas, B.S., et al., The locus coeruleus drives disinhibition in the midline thalamus via a dopaminergic mechanism. Nat Neurosci, 2018. 21(7): p. 963–973.

62. Zhang, L., M. Kolaj, and L.P. Renaud, Intracellular postsynaptic cannabinoid receptors link thyrotropin-releasing hormone receptors to TRPC-like channels in thalamic paraventricular nucleus neurons. Neuroscience, 2015. 311: p. 81–91.

63. Zhang, L., M. Kolaj, and L.P. Renaud, Endocannabinoid 2-AG and intracellular cannabinoid receptors modulate a low-threshold calcium spike-induced slow depolarizing afterpotential in rat thalamic paraventricular nucleus neurons. Neuroscience, 2016. 322: p. 308–319.

64. Kolaj, M., et al., Intrinsic properties and neuropharmacology of midline paraventricular thalamic nucleus neurons. Front Behav Neurosci, 2014. 8: p. 132.

65. Zhang, L., L.P. Renaud, and M. Kolaj, Properties of a T-type Ca2+channel-activated slow afterhyperpolarization in thalamic paraventricular nucleus and other thalamic midline neurons. J Neurophysiol, 2009. 101(6): p. 2741–50.

66. Zhang, L., M. Kolaj, and L.P. Renaud, Ca2+-dependent and Na+-dependent K+ conductances contribute to a slow AHP in thalamic paraventricular nucleus neurons: a novel target for orexin receptors. J Neurophysiol, 2010. 104(4): p. 2052–62.

67. Richter, T.A., M. Kolaj, and L.P. Renaud, Low voltage-activated Ca2+ channels are coupled to Ca2+-induced Ca2+ release in rat thalamic midline neurons. J Neurosci, 2005. 25(36): p. 8267–71.

68. Jones, E.G. and S.H. Hendry, Differential Calcium Binding Protein Immunoreactivity Distinguishes Classes of Relay Neurons in Monkey Thalamic Nuclei. Eur J Neurosci, 1989. 1(3): p. 222–246.

69. Jones, E.G., A new view of specific and nonspecific thalamocortical connections. Adv Neurol, 1998. 77: p. 49–71; discussion 72-3.

70. Jones, E.G., Viewpoint: the core and matrix of thalamic organization. Neuroscience, 1998. 85(2): p. 331–45.

71. Jones, E.G., The thalamic matrix and thalamocortical synchrony. Trends in Neurosciences, 2001. 24(10): p. 595–601.

72. Sherman, S.M. and R.W. Guillery, The role of the thalamus in the flow of information to the cortex. Philos Trans R Soc Lond B Biol Sci, 2002. 357(1428): p. 1695-708.

73. Clascá, F., P. Rubio-Garrido, and D. Jabaudon, Unveiling the diversity of thalamocortical neuron subtypes. Eur J Neurosci, 2012. 35(10): p. 1524–32.

74. Halassa, M.M. and S.M. Sherman, Thalamocortical Circuit Motifs: A General Framework. Neuron, 2019. 103(5): p. 762–770.

75. Li, S. and G.J. Kirouac, Sources of inputs to the anterior and posterior aspects of the paraventricular nucleus of the thalamus. Brain Struct Funct, 2012. 217(2): p. 257–73.

76. Zingg, B., et al., Neural networks of the mouse neocortex. Cell, 2014. 156(5): p. 1096–111.

77. Su, H.S. and M. Bentivoglio, Thalamic midline cell populations projecting to the nucleus accumbens, amygdala, and hippocampus in the rat. J Comp Neurol, 1990. 297(4): p. 582–93.

78. Campus, P., et al., The paraventricular thalamus is a critical mediator of top-down control of cue-motivated behavior in rats. Elife, 2019. 8.

79. Otis, J.M., et al., Paraventricular thalamus projection neurons integrate cortical and hypothalamic signals for cue-reward processing. Neuron, 2019. 102(5).

80. Do-Monte, F.H., K. Quinones-Laracuente, and G.J. Quirk, A temporal shift in the circuits mediating retrieval of fear memory. Nature, 2015. 519(7544): p. 460-3.

81. Quiñones-Laracuente, K., A. Vega-Medina, and G.J. Quirk, Time-Dependent Recruitment of Prelimbic Prefrontal Circuits for Retrieval of Fear Memory. Frontiers in Behavioral Neuroscience, 2021. 15.

82. Otis, J.M., et al., Prefrontal cortex output circuits guide reward seeking through divergent cue encoding. Nature, 2017. 543(7643): p. 103-107.

83. Haight, J.L. and S.B. Flagel, A potential role for the paraventricular nucleus of the thalamus in mediating individual variation in Pavlovian conditioned responses. Front Behav Neurosci, 2014. 8: p. 79.

84. Cain, C.K. and J.E. LeDoux, Chapter 3.1 Brain mechanisms of Pavlovian and instrumental aversive conditioning, in Handbook of Behavioral Neuroscience, R.J. Blanchard, et al., Editors. 2008, Elsevier. p. 103–124.

85. Martinez, R.C.R., et al., Active vs. reactive threat responding is associated with differential c-Fos expression in specific regions of amygdala and prefrontal cortex. Learning & memory (Cold Spring Harbor, N.Y.), 2013. 20(8): p. 446–452.

86. Penzo, M.A., et al., The paraventricular thalamus controls a central amygdala fear circuit. Nature, 2015. 519(7544): p. 455-9.

87. Li, Y., et al., Lesions of the posterior paraventricular nucleus of the thalamus attenuate fear expression. Frontiers in Behavioral Neuroscience, 2014. 8(94).

88. Ma, J., et al., Divergent projections of the paraventricular nucleus of the thalamus mediate the selection of passive and active defensive behaviors. Nat Neurosci, 2021. 24(10): p. 1429–1440.

89. Lee, H., et al., Scalable control of mounting and attack by Esr1+ neurons in the ventromedial hypothalamus. Nature, 2014. 509(7502): p. 627-632.

90. Hashikawa, K., et al., Esr1(+) cells in the ventromedial hypothalamus control female aggression. Nat Neurosci, 2017. 20(11): p. 1580–1590.

91. Falkner, A.L. and D. Lin, Recent advances in understanding the role of the hypothalamic circuit during aggression. Frontiers in Systems Neuroscience, 2014. 8.

92. Liu, M., et al., Make war not love: The neural substrate underlying a state-dependent switch in female social behavior. Neuron, 2022. 110(5): p. 841–856.e6.

93. Lo, L., et al., Connectional architecture of a mouse hypothalamic circuit node controlling social behavior. Proceedings of the National Academy of Sciences, 2019. 116(15): p. 7503–7512.

94. Cheng, Y.-F., et al., Cardioprotection induced in a mouse model of neuropathic pain via anterior nucleus of paraventricular thalamus. Nature Communications, 2017. 8(1): p. 826.

95. Do-Monte, F.H., et al., Thalamic Regulation of Sucrose Seeking during Unexpected Reward Omission. Neuron, 2017. 94(2): p. 388–400 e4.

96. Lähnemann, D., et al., Eleven grand challenges in single-cell data science. Genome Biology, 2020. 21(1): p. 31.

97. Lafferty, C.K., et al., Nucleus Accumbens Cell Type- and Input-Specific Suppression of Unproductive Reward Seeking. Cell Reports, 2020. 30(11): p. 3729–3742.e3.

98. Huang, H., P. Ghosh, and A.N. van den Pol, Prefrontal Cortex–Projecting Glutamatergic Thalamic Paraventricular Nucleus-Excited by Hypocretin: A Feedforward Circuit That May Enhance Cognitive Arousal. Journal of Neurophysiology, 2006. 95(3): p. 1656–1668.

99. Matzeu, A., E. Zamora-Martinez, and R. Martin-Fardon, The paraventricular nucleus of the thalamus is recruited by both natural rewards and drugs of abuse: recent evidence of a pivotal role for orexin/hypocretin signaling in this thalamic nucleus in drug-seeking behavior. Frontiers in Behavioral Neuroscience, 2014. 8(117).

100. Matzeu, A. and R. Martin-Fardon, Drug Seeking and Relapse: New Evidence of a Role for Orexin and Dynorphin Co-transmission in the Paraventricular Nucleus of the Thalamus. Front Neurol, 2018. 9: p. 720.

101. Li, Y., et al., Changes in emotional behavior produced by orexin microinjections in the paraventricular nucleus of the thalamus. Pharmacol Biochem Behav, 2010. 95(1): p. 121–8.

102. Li, Y., et al., Orexins in the paraventricular nucleus of the thalamus mediate anxiety-like responses in rats. Psychopharmacology (Berl), 2010. 212(2): p. 251–65.

103. Choi, D.L., et al., Orexin signaling in the paraventricular thalamic nucleus modulates mesolimbic dopamine and hedonic feeding in the rat. Neuroscience, 2012. 210: p. 243–8.

104. Cole, S., H.S. Mayer, and G.D. Petrovich, Orexin/Hypocretin-1 Receptor Antagonism Selectively Reduces Cue-Induced Feeding in Sated Rats and Recruits Medial Prefrontal Cortex and Thalamus. Sci Rep, 2015. 5: p. 16143.

105. Heydendael, W., et al., Orexins/hypocretins act in the posterior paraventricular thalamic nucleus during repeated stress to regulate facilitation to novel stress. Endocrinology, 2011. 152(12): p. 4738–52.

106. Heydendael, W., A. Sengupta, and S. Bhatnagar, Putative genes mediating the effects of orexins in the posterior paraventricular thalamus on neuroendocrine and behavioral adaptations to repeated stress. Brain Res Bull, 2012. 89(5-6): p. 203–10.

107. Choi, D.L., et al., The role of orexin-A in food motivation, reward-based feeding behavior and food-induced neuronal activation in rats. Neuroscience, 2010. 167(1): p. 11–20.

108. Dong, X., Y. Li, and G.J. Kirouac, Blocking of orexin receptors in the paraventricular nucleus of the thalamus has no effect on the expression of conditioned fear in rats. Frontiers in Behavioral Neuroscience, 2015. 9.

109. Ren, S., et al., The paraventricular thalamus is a critical thalamic area for wakefulness. Science, 2018. 362(6413): p. 429-434.

110. Sharma, S., C. Hryhorczuk, and S. Fulton, Progressive-ratio responding for palatable high-fat and high-sugar food in mice. Journal of visualized experiments: JoVE, 2012(63): p. e3754–e3754.

111. Lee, Y.H., et al., Food Craving, Seeking, and Consumption Behaviors: Conceptual Phases and Assessment Methods Used in Animal and Human Studies. J Obes Metab Syndr, 2019. 28(3): p. 148–157.

112. Dalmay, T., et al., A Critical Role for Neocortical Processing of Threat Memory. Neuron, 2019. 104(6): p. 1180–1194.e7.

113. Cho, J.-H., B.S. Huang, and J.M. Gray, RNA sequencing from neural ensembles activated during fear conditioning in the mouse temporal association cortex. Scientific Reports, 2016. 6(1): p. 31753.

114. Yu, H.H., et al., A specialized area in limbic cortex for fast analysis of peripheral vision. Curr Biol, 2012. 22(14): p. 1351–7.

115. Haight, J.L., et al., A food-predictive cue attributed with incentive salience engages subcortical afferents and efferents of the paraventricular nucleus of the thalamus. Neuroscience, 2017. 340: p. 135–152.

116. Zhu, Y., et al., Dynamic salience processing in paraventricular thalamus gates associative learning. Science, 2018. 362(6413): p. 423-429.

117. Garau, C., et al., Neuropeptide S Encodes Stimulus Salience in the Paraventricular Thalamus. Neuroscience, 2022.

118. Deutch, A.Y., M. Bubser, and C.D. Young, Psychostimulant-induced Fos protein expression in the thalamic paraventricular nucleus. J Neurosci, 1998. 18(24): p. 10680–7.

119. Levant, B., D.E. Grigoriadis, and E.B. DeSOUZA, [3H] quinpirole binding to putative D2 and D3 dopamine receptors in rat brain and pituitary gland: a quantitative autoradiographic study. Journal of Pharmacology and Experimental Therapeutics, 1993. 264(2): p. 991–1001.

120. Ao, Y., et al., Application of quinpirole in the paraventricular thalamus facilitates emergence from isoflurane anesthesia in mice. Brain and Behavior, 2020. **n/a**(n/a): p. e01903.

121. Ao, Y., et al., Locus Coeruleus to Paraventricular Thalamus Projections Facilitate Emergence From Isoflurane Anesthesia in Mice. Front Pharmacol, 2021. 12: p. 643172.

122. Davis, M.I., et al., The cannabinoid-1 receptor is abundantly expressed in striatal striosomes and striosome-dendron bouquets of the substantia nigra. PloS one, 2018. 13(2): p. e0191436–e0191436.

123. Krout, K.E., R.E. Belzer, and A.D. Loewy, Brainstem projections to midline and intralaminar thalamic nuclei of the rat. J Comp Neurol, 2002. 448(1): p. 53–101.

124. Vertes, R.P., A PHA-L analysis of ascending projections of the dorsal raphe nucleus in the rat. J Comp Neurol, 1991. 313(4): p. 643–68.

125. Krout, K.E. and A.D. Loewy, Periaqueductal gray matter projections to midline and intralaminar thalamic nuclei of the rat. J Comp Neurol, 2000. 424(1): p. 111–41.

126. Zhang, L., et al., Vasopressin induces depolarization and state-dependent firing patterns in rat thalamic paraventricular nucleus neurons in vitro. Am J Physiol Regul Integr Comp Physiol, 2006. 290(5): p. R1226–32.

127. Zhang, L., M. Kolaj, and L.P. Renaud, GIRK-like and TRPC-like conductances mediate thyrotropin-releasing hormone-induced increases in excitability in thalamic paraventricular nucleus neurons. Neuropharmacology, 2013. 72: p. 106–15.

128. Kolaj, M., et al., Orexin-induced modulation of state-dependent intrinsic properties in thalamic paraventricular nucleus neurons attenuates action potential patterning and frequency. Neuroscience, 2007. 147(4): p. 1066–75.

129. Ishibashi, M., et al., Effects of orexins/hypocretins on neuronal activity in the paraventricular nucleus of the thalamus in rats in vitro. Peptides, 2005. 26(3): p. 471–81.

130. Ong, Z.Y., et al., Paraventricular Thalamic Control of Food Intake and Reward: Role of Glucagon-Like Peptide-1 Receptor Signaling. Neuropsychopharmacology, 2017. 42(12): p. 2387–2397.

131. Heilbronner, U., M. van Kampen, and G. Flügge, The Alpha-2B Adrenoceptor in the Paraventricular Thalamic Nucleus is Persistently Upregulated by Chronic Psychosocial Stress. Cellular and Molecular Neurobiology, 2004. 24(6): p. 815.

132. Kark, S.M., et al., Functional Connectivity of the Human Paraventricular Thalamic Nucleus: Insights From High Field Functional MRI. Frontiers in Integrative Neuroscience, 2021. 15.

133. Rieck, R.W., et al., Distribution of dopamine D2-like receptors in the human thalamus: autoradiographic and PET studies. Neuropsychopharmacology, 2004. 29(2): p. 362–72.

134. Nagalski, A., et al., Molecular anatomy of the thalamic complex and the underlying transcription factors. Brain Struct Funct, 2016. 221(5): p. 2493–510.

135. Kelley, A.E., B.A. Baldo, and W.E. Pratt, A proposed hypothalamic–thalamic– striatal axis for the integration of energy balance, arousal, and food reward. Journal of Comparative Neurology, 2005. 493(1): p. 72–85.

136. Zheng, G.X.Y., et al., Massively parallel digital transcriptional profiling of single cells. Nature Communications, 2017. 8(1): p. 14049.

137. Russ, D.E., et al., A harmonized atlas of mouse spinal cord cell types and their spatial organization. Nature Communications, 2021. 12(1): p. 5722.

138. Saunders, A., et al., Molecular Diversity and Specializations among the Cells of the Adult Mouse Brain. Cell, 2018. 174(4): p. 1015–1030.e16.

139. Erben, L., et al., A Novel Ultrasensitive In Situ Hybridization Approach to Detect Short Sequences and Splice Variants with Cellular Resolution. Molecular neurobiology, 2018. 55(7): p. 6169–6181.

140. Erben, L. and A. Buonanno, Detection and Quantification of Multiple RNA Sequences Using Emerging Ultrasensitive Fluorescent In Situ Hybridization Techniques. Current protocols in neuroscience, 2019. 87(1): p. e63.

